# Macrophage renewal in the small intestine governs early-life control of streptococci

**DOI:** 10.64898/2026.02.11.705249

**Authors:** Zohreh Mansoori Moghadam, Mirjam Freudenhammer, Maria Francesca Viola, Marleen Eckert, Candice Raynaud, Linus Kriegl, Michael Rauer, Reem Alsumati, Nicole Treichel, Ramona Rebekka Eckert, Sebastian Baasch, Melanie Boerries, Elvira Mass, Thomas Clavel, Daniel Erny, Julia Kolter, Philipp Henneke

## Abstract

Group B Streptococcus (GBS) is both a common intestinal colonizer that is transmitted intergenerationally, and a primary cause of neonatal sepsis worldwide. However, the innate immune mechanisms that restrict the pathogen at the intestinal barrier early in life remain poorly understood. Using an enteral GBS-colonization model in infant mice, we found that lamina-propria (LP) macrophages controlled both GBS-colonization densities and invasion in an age-dependent fashion. LP macrophages turnover and differentiation were strongly impacted by topology. In the small intestine, monocytes infiltration of the LP occurred from birth on in response to perinatally acquired microbiota, whereas in the colon it was driven by the weaning reaction. Moreover, macrophages of the small intestine mounted a robust, MyD88-dependent inflammatory response to GBS, while those of the colon remained largely unaffected. Together, these findings demonstrate that region-specific macrophage dynamics early in life critically influence host–pathogen-interactions during GBS-colonization.

## Introduction

Group B Streptococcus (GBS) is a leading cause of neonatal sepsis and meningitis worldwide. Clinically, neonatal GBS disease can be divided in an early-onset disease (EOD) occurring within the first week of life and a late-onset disease (LOD) that develops from day 7 until day 90 after birth ^1^. In EOD, infection with GBS is thought to occur during birth by inhalation of infected vaginal fluids. Accordingly, EOD is substantially reduced by intrapartum antibiotic prophylaxis ^2^. In LOD, GBS infection probably arises after GBS colonization of the intestine and subsequent translocation across mucosal barriers ^3,4^. However, the mechanisms that promote invasive disease rather than asymptomatic colonization - as is the case in the majority of infants - remain unclear ^5,6^.

Neonates exhibit a strong inflammatory cytokine response to GBS but simultaneously display impaired bacterial clearance, suggesting that developmental immune factors may contribute to their heightened susceptibility to GBS infection ^7,8^. Macrophages play a central role among intestinal immune cells, maintaining gut homeostasis and promoting wound healing and epithelial repair ^9,10^. In the adult colon the majority of macrophages are prenatally seeded cells with limited life span and are continuously replenished from circulating blood monocytes that enter the mucosa and differentiate. This process is shaped by local cues ^11^. Recruitment of classical CCR2⁺Ly6C^high^ monocytes is a major pathway for replenishment in steady state and during inflammation, although subsets of more self-maintaining, foetal-derived macrophages also exist in defined intestinal niches ^12,13^. Germ-free (GF) mice show impaired macrophage maturation and altered monocyte-to-macrophage turnover in the gut, and conventional microbiota colonization restores normal recruitment, maturation and turnover kinetics ^14,15^. The majority of intestinal resident macrophages are localized within the lamina propria (LP) of the mucosa, where they are crucial for the defense against invading microbes and for the interaction with dietary antigens ^12,16^. Consequently, LP macrophages are central to maintaining the balance between microbial defense and oral tolerance ^17^. However, their role in restraining invasive pathogens such as GBS at the intestinal interface during early life remains poorly understood.

To address this gap, we developed and characterized a murine model of LOD. Using oral inoculation of mice at different developmental stages, we observed that intestinal colonization by GBS was age dependent. Furthermore, small intestinal macrophages from neonatal mice displayed heightened responsiveness to GBS compared to colonic macrophages, suggesting compartment-specific regulation that may influence outcome after natural early-life exposure to a pathogen.

## Results

### GBS colonization depends on age and macrophage abundance

To investigate the influence of age on host susceptibility to GBS, a murine GBS colonization model was established in which C57BL/6 mice of different ages were orally inoculated with GBS **(Figure 1A)**. Intestinal colonization by GBS exhibited a clear age dependence as neonatal mice acquired colonization more readily and maintained bacterial carriage for a longer duration compared to adult mice **(Figure 1B)**. Despite these differences in colonization dynamics, colonized wild-type (WT) mice of all ages survived for at least 21 days post-colonization **(Figure 1C)**. In the intestine, Ly6C⁺MHC-II⁻ monocytes regularly migrate into the LP, where they differentiate into Ly6C⁻MHC-II⁺ tissue-resident macrophages **(Figure S1A)**. Given the important role of intestinal macrophages in maintaining gut homeostasis ^12^, we hypothesized that they played a critical role in age-dependent susceptibility to GBS. To test this, macrophages were depleted with an orally administrated colony-stimulating factor 1 receptor (CSF1R) inhibitor (BSL) **(Figure 1D).** Efficient depletion of intestinal macrophages was verified by both flow cytometry, and immunofluorescence analyses **(Figure 1E, F, and S1B-C).** Following macrophage depletion, we found significantly lower survival rates **(Figure 1G)**, in addition to higher GBS loads in fecal samples **(Figure 1H),** and strongly increased rates of dissemination to peripheral tissues, including the liver, spleen, and mesenteric lymph nodes, at 2- and 4-days post- colonization **(Figure 1I, J)**. This indicated a strong protective role of macrophages against both GBS colonization and invasion. As the gut macrophage compartment is rapidly reconstituted within days due to the recruitment and differentiation of circulating CCR2⁺ Ly6C⁺ monocytes (Bain et al. 2014). We next analyzed *Ccr2^⁻/⁻^* mice, in which monocyte egress from the bone marrow and recruitment to peripheral tissues are markedly impaired ^18,19^. The combined approach, genetic disruption of CCR2 alongside CSF1R inhibition, thus establishes an extended and more complete window for investigating macrophage-dependent processes during GBS colonization **(Figure 1K)**. Strikingly, this dual intervention resulted in an even more pronounced, i.e. faster reduction in survival, with approximately 40% of *Ccr2^⁻/⁻^* mice dying within two days post-colonization **(Figure 1L)**. Analysis of fecal samples revealed elevated GBS loads in *Ccr2^⁻/⁻^* mice compared to wild-type controls under BSL treatment **(Figure 1M)**. Furthermore, GBS dissemination occurred more frequently and extensively in *Ccr2^⁻/⁻^* mice treated with BSL, indicating a critical role for monocyte-derived macrophages in restricting bacterial dissemination **(Figure 1N)**. Collectively, these findings demonstrated that deficiency of intestinal monocytes and macrophages compromised the host’s ability to control intestinal GBS colonization and dissemination. This indicated an essential contribution of monocyte-derived macrophages to mucosal antibacterial defense, especially in the neonate.

**Figure 1.**
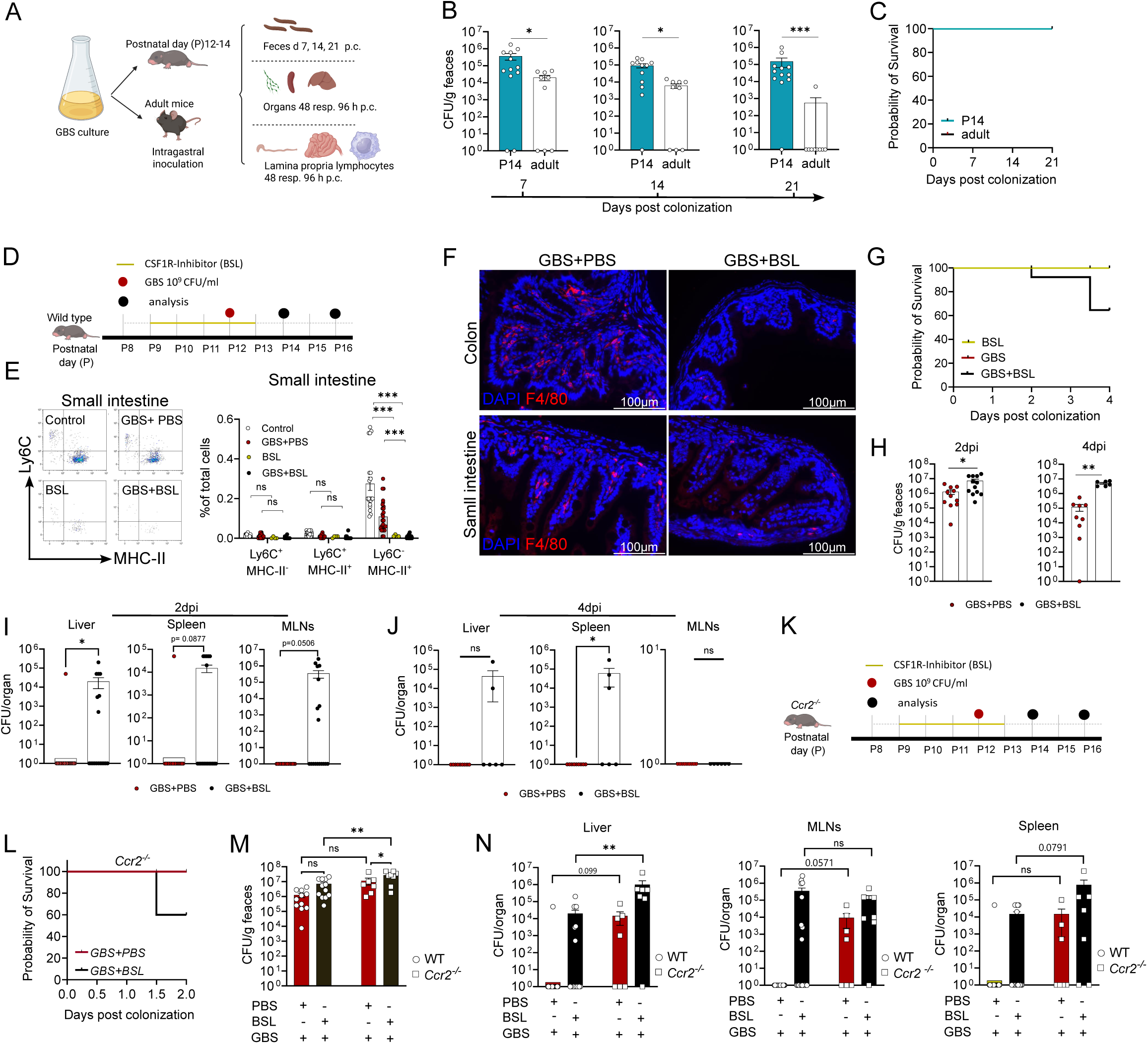
GBS colonization depends on age and macrophage abundance. **A.** Schematic of the experimental plan. Neonatal mice were infected between P12–P14 with 10^9^ colony forming unit (CFU)/ml GBS via oral gavage. Adult mice were also colonized with 10^10^ CFU/ml GBS. **B.** Quantification of Log₁₀ colony-forming units (CFU) per gram of feces from wild-type (WT) C57BL/6 mice at 7-, 14-, and 21-days post GBS colonization. Data represents P14 (n = 12) and adult (n = 9) mice, combined from two independent experiments. **C.** Survival rate following GBS colonization at postnatal day 14 and adult mice 21 days post colonization. Data represents P14 (n = 12) and adult (n = 9) mice, combined from two independent experiments. **D.** Schematic representation of experimental design. WT mice received daily administration of the CSF1 receptor inhibitor (BSL) for five consecutive days, starting at postnatal day 9 (P9). Bacterial colonization was performed at P12. Experimental analyses were conducted either 2- or 4-days post-colonization. **E.** Flow cytometric assessment of macrophage depletion efficiency in the small intestine. Tissue-resident macrophages were identified by sequential gating on CD45⁺ CD11b⁺ Ly6G⁻ SiglecF⁻ CD64⁺ cells, 2 days post colonization. Experimental groups included Control (n = 20), GBS+PBS (n= 27), BSL (n= 4), and BSL+GBS (n=15), analyzed across 4 independent experiments. **F.** Evaluation of depletion efficiency in the colon and small intestine by immunofluorescence for F4/80⁺ macrophages at 2 days post colonization. Scale bar indicates 100µm. **G.** Survival rate of WT mice following GBS colonization in addition to the BSL treatment at 4 days post colonization, data represent BSL (n= 4), GBS (n= 19), and GBS+BSL (n= 18) mice. **H.** Quantification of Log₁₀ CFU of the GBS per gram of feces from WT C57BL/6 mice at 2-, and 4-days post GBS colonization. Data represent GBS+PBS (n= 18) and GBS+BSL (n= 18) mice, combined from 3 independent experiments. **I.** Quantification of GBS dissemination to the liver, spleen, and mesenteric lymph nodes (MLNs) 2 days post-colonization. Data represents combined results from two independent experiments, including GBS+PBS (n= 18) and GBS+BSL (n= 18) mice. **J.** Quantification of GBS dissemination to the liver, spleen, and mesenteric lymph nodes (MLNs) 4 days post-colonization. Data represents combined results from two independent experiments, including GBS+PBS (n= 18) and GBS+BSL (n= 18) mice, combined from 3 independent experiments. **K.** Schematic representation of experimental design. *Ccr2^-/-^*mice received daily administrations of the CSF1 receptor inhibitor (BSL) for five consecutive days, starting at postnatal day 9 (P9). Bacterial colonization was performed at P12. Experimental analyses were conducted 2 days post-colonization. **L.** Survival rate of *Ccr2^-/-^* mice following GBS colonization in addition to the BSL treatment at 2 days post colonization, GBS+PBS (n= 8) and GBS+BSL (n= 10) mice, in 2 individual experiments. **M.** Quantification of Log₁₀ CFU of the GBS per gram of feces from *Ccr2^-/-^* mice at 2-days post GBS colonization. Data represent in GBS+PBS (n= 18) and GBS+BSL (n= 18) WT mice, and with *Ccr2^-/-^* mice GBS+PBS (n= 8) and GBS+BSL (n= 7) combined from 3 independent experiments. **N.** Quantification of GBS dissemination to the liver, spleen, and mesenteric lymph nodes (MLNs) 2 days post-colonization. Data represent in GBS+PBS (n= 18) and GBS+BSL (n= 18) WT mice, and with *Ccr2^-/-^* mice GBS+PBS (n= 8) and GBS+BSL (n= 7) combined from 3 independent experiments.

### Distinct developmental programs of small intestine and colonic macrophages

The dependence of GBS colonization on age and macrophage abundance suggested that intestinal macrophages undergo dynamic changes during development. The small and large intestine follow distinct developmental programs during embryogenesis. Villi in the small intestine start to form already around E14.5 ^20^, whereas morphological maturation of the colon occurs mostly postnatally ^21^. Despite the different morphogenetic developmental patterns of small intestine and colon, the immunophenotypic trajectories of monocytes and macrophages, are thought to be largely overlapping^16,22^. However, most studies have focused on colon macrophages and functional data on small intestine macrophages remain incomplete ^11,22–24^.

To investigate region-specific macrophage dynamics in the gut-associated compartments, we analyzed developmental time points from embryonic day (E)18.5 to postnatal day (P)42 (E18.5, P1, P3, P8, P12, P21, and P42) **(Figure 2A)**. During early development (E18.5 and P3), we observed a significant increase in Ly6C⁺MHC-II⁻ monocytes in both the colon and small intestine. At the time of weaning (P21), monocyte numbers were significantly elevated only in the colon, suggesting that the response to the weaning transition ^11^ was colon-specific **(Figure 2B)**. Consistently, the frequency of the Ly6C⁺MHC-II⁺ macrophage population also increased exclusively in the colon at P21 **(Figure 2C).** As confirmation of a prominent weaning reaction occurring in the colon compared to the small intestine at P21, we observed a significant upregulation of the proinflammatory cytokine *Tnf* and the chemotactic gene *Cxcl1*, primarily in colon Ly6C⁺MHC-II⁺ macrophages **(Figure 2D).** Notably, at E18.5, the small intestine harbored a substantially higher proportion of tissue-resident Ly6C⁻MHC-II⁺ macrophages which progressively declined during postnatal development (**Figure 2E)**. Immunofluorescence analysis of E18.5 tissues demonstrated that MHC-II⁺ macrophages were abundantly localized within the LP of the small intestine. Conversely, in the colon, MHC-II⁺ macrophages were only infrequently detected in the laminar regions, indicating tissue-specific differences in macrophage distribution at this developmental stage **(Figure 2F).** This was in line with the published lack of MHC-II expression in colon macrophages in early life (Bain et al, 2014). Ly6C⁻MHC-II⁻ macrophages exhibited prolonged persistence in the colon, whereas they largely disappeared in the small intestine by P1, representing a notable spatial and temporal difference in macrophage dynamics **(Figure 2G)**. Accordingly, the mean fluorescence intensity (MFI) of MHC-II in colon macrophages gradually increased from E18.5 to adulthood **(Figure 2H)**, while it reached adult levels already at E18.5 in the small intestine and then showed minimal variation over time **(Figure 2I)**. Interestingly, CD206⁺ macrophages, which constitute approximately 50% of adult colonic macrophages, appeared to maintain a similar proportion from before birth through adulthood **(Figure 2J)**. However, in the small intestine, CD206 was strongly upregulated immediately after birth in almost all small intestinal macrophages during the early postnatal period (around P1–P3), and then decreased to 40% of the population **(Figure 2K)**. Immunofluorescence staining of the colon and small intestine in mice at P3 were in line with data generated by flow cytometry **(Figure 2L)**. This indicated that the majority of small intestinal macrophages were CD206⁺ during early postnatal life. Elevated expression of CD206 by macrophages in the neonatal small intestine may suggest a role for these cells in early postnatal immune adaptation, possibly contributing to tissue remodeling and the establishment of immune homeostasis associated ^25,26^, with the critical transition from fetal to adult life. Taken together and considering that the morphogenesis of the small intestine precedes that of the large intestine, these findings suggest that mononuclear phagocytes in this region are primed substantially earlier in life and may play a pivotal role in early immune defense against GBS colonization.

**Figure 2.**
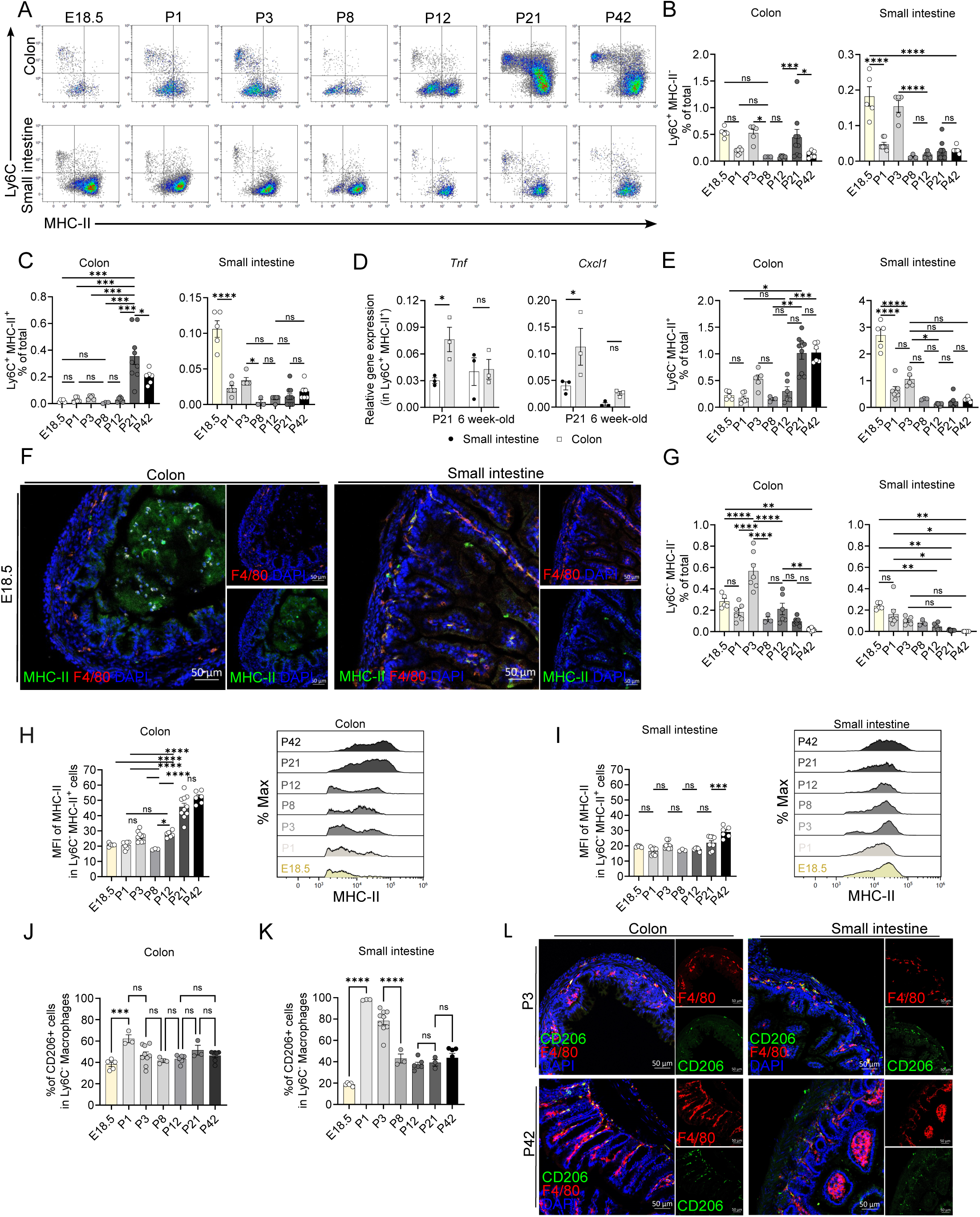
Distinct developmental programs of small intestine and colonic macrophages. **A.** Flow cytometric analysis of Ly6C and MHC-II expression in CD45⁺CD11b⁺SiglecF⁻Ly6G⁻CD64⁺ macrophages isolated from the lamina propria (LP) of the colon and small intestine at critical developmental stages (embryonic day 18.5 (E18.5), postnatal day 1 (P1), P3, P8, P12, P21, P42. **B.** Quantification of Ly6C⁺MHC-II⁻ monocytes in the colon and small intestine at E18.5 (n= 5) and P1 (n= 3), P3 (n= 6), P8 (n= 3), P12(n= 6), P21(n= 6), P42 (n=7) as determined by flow cytometry. **C.** Quantification of Ly6C⁺MHC-II^+^ monocytes-derived macrophages in the colon and small intestine at E18.5 (n= 5) and P1 (n= 3), P3 (n= 6), P8 (n= 3), P12(n= 6), P21(n= 6), P42 (n=7) as determined by flow cytometry. **D.** Quantitative real-time PCR (qRT-PCR) analysis of *Tnfa* and *Cxcl1* gene expression in Ly6C⁺ MHC-II⁺ cells isolated from the colon and small intestine of p21 (weaning age) and adult (6-week-old) mice. **E.** Quantification of Ly6C^-^MHC-II^+^ macrophages in the colon and small intestine at E18.5 (n= 5) and P1 (n= 3), P3 (n= 6), P8(n= 3), P12(n= 6), P21(n= 6), P42 (n= 7) as determined by flow cytometry. **F.** Immunofluorescence staining of MHC-II (green), F4/80 (red), and DAPI (blue) in distal colon and distal small intestine of E18.5 embryos. Scale bar: 50 µm. **G.** Quantification of Ly6C^-^MHC-II⁻ macrophages in the colon and small intestine at E18.5 (n= 5) and P1 (n= 3), P3 (n= 6), P8 (n= 3), P12 (n= 6), P21 (n= 6), P42 (n=7) as determined by flow cytometry. **H.** Quantitative analysis of the mean fluorescence intensity (MFI) of MHC-II expression in colonic Ly6C⁻MHC-II⁺ macrophages at E18.5 and postnatal days 1, 3, 8, 12, 21, and 42 (P1, P3, P8, P12, P21, and P42). **I.** Quantitative analysis of the MFI of MHC-II expression in small intestinal Ly6C⁻MHC-II⁺ macrophages at E18.5, P1, P3, P8, P12, P21, and P42. **J.** Quantitative analysis of the MFI of CD206 expression in small intestinal Ly6C⁻MHC-II⁺ macrophages at E18.5, P1, P3, P8, P12, P21, and P42. **K.** Quantitative analysis of the MFI of CD206 expression in small intestinal Ly6C⁻MHC-II⁺ macrophages at E18.5, P1, P3, P8, P12, P21, and P42. **L.** Immunofluorescence staining of CD206 (green), F4/80 (red), and DAPI (blue) in distal colon and distal small intestine of P3, and P42 mice. Scale bar: 50 µm.

### Distinct ontogenic dynamics of tissue-resident macrophages in small intestine and colon

The development of tissue-resident macrophages in the adult intestine is relatively well characterized ^10^, however, their ontogeny during prenatal and neonatal stages, particularly in the small intestine, remains poorly understood. To investigate the developmental origin of colonic and small intestinal macrophages during development, we utilized a double fate-mapping model, *Tnfrsf11a^Cre^; Rosa26^LSL-YFP^; Ms4a3^FlpO^; Rosa26^FSF-tdTomato^* ^27^, which enables the simultaneous tracing of yolk sac-derived and bone marrow-derived cells. In this model, yolk sac pre-macrophages express *Tnfrsf11a* are permanently labeled with YFP, resulting in YFP expression in all tissue-resident macrophages derived from this lineage^28^, while *Ms4a3* labels monocyte-derived macrophages originating from the granulocyte-monocyte progenitor (GMP) lineage with tdTomato ^29^ **(Figure 3A)**. Notably, *Tnfrsf11a* is a macrophage core gene that becomes upregulated in monocyte-derived macrophages once they become resident. Using this model, we performed flow cytometric analyses of the colon and small intestine across developmental stages. In line with their bone marrow origin, Ly6C⁺MHC-II⁻ monocytes were predominantly tdTomato^+^ and YFP^-^ **(Figure 3B, C)**. Interestingly, the proportion of tdTomato⁺ Ly6C⁺MHC-II⁺ macrophages progressively increased in the colon, whereas in the small intestine it remained relatively stable until P21, after which it began to rise **(Figure 3D)**. In contrast, *Tnfrsf11a* expression, and consequently YFP labelling, was upregulated earlier in the small intestine than in the colon prior to weaning **(Figure 3E)**, indicating earlier macrophage establishment and/or longer tissue residency in the small intestine. For mature Ly6C⁻MHC-II⁺macrophages, our data demonstrated that the weaning-associated shift to bone marrow-derived macrophages again occurred primarily in the colon, as evidenced by a significant increase in tdTomato*^+^* macrophages within this compartment at P21. In the small intestine, on the other hand YFP⁺ macrophages were maintained and a slow influx of bone marrow-derived cells occurred predominantly in the adult age **(Figure 3F, G)**. Notably, both compartments showed tdTomato labeling of approximately 50% of macrophages in the adult, in line with only a partial replacement of embryonic macrophages ^29,30^. Immunofluorescence staining further corroborated the flow cytometry data. At P21, most tissue-resident macrophages in the colon were tdTomato⁺ (i.e. Ms4a3+), confirming the progressive recruitment of bone marrow–derived macrophages during the weaning period. In contrast, this replacement was minimal in the small intestine, indicating tissue-specific regulation of macrophage turnover **(Figure 3H)**. Moreover, in the colon, the majority of Ly6C⁻MHC-II⁻ macrophages did not express tdTomato prior to weaning, suggesting that these cells were largely not GMP-derived **(Figure 3I)**. This observation is consistent with the notion that Ly6C⁻MHC-II⁻ colonic macrophages are predominantly of embryonic origin, by contrast to the small intestine **(Figure 3J)**. Together, these findings demonstrate that the contribution of bone marrow–derived monocytes to the macrophage pool in the gut is region-dependent and developmentally regulated. The colon undergoes extensive postnatal remodeling of its macrophage compartment around weaning, characterized by increased recruitment and integration of monocyte-derived macrophages. Conversely, the small intestine maintains a more stable population of pre-macrophage-derived macrophages, suggesting distinct mechanisms govern macrophage ontogeny and maintenance in these different tissue environments.

**Figure 3.**
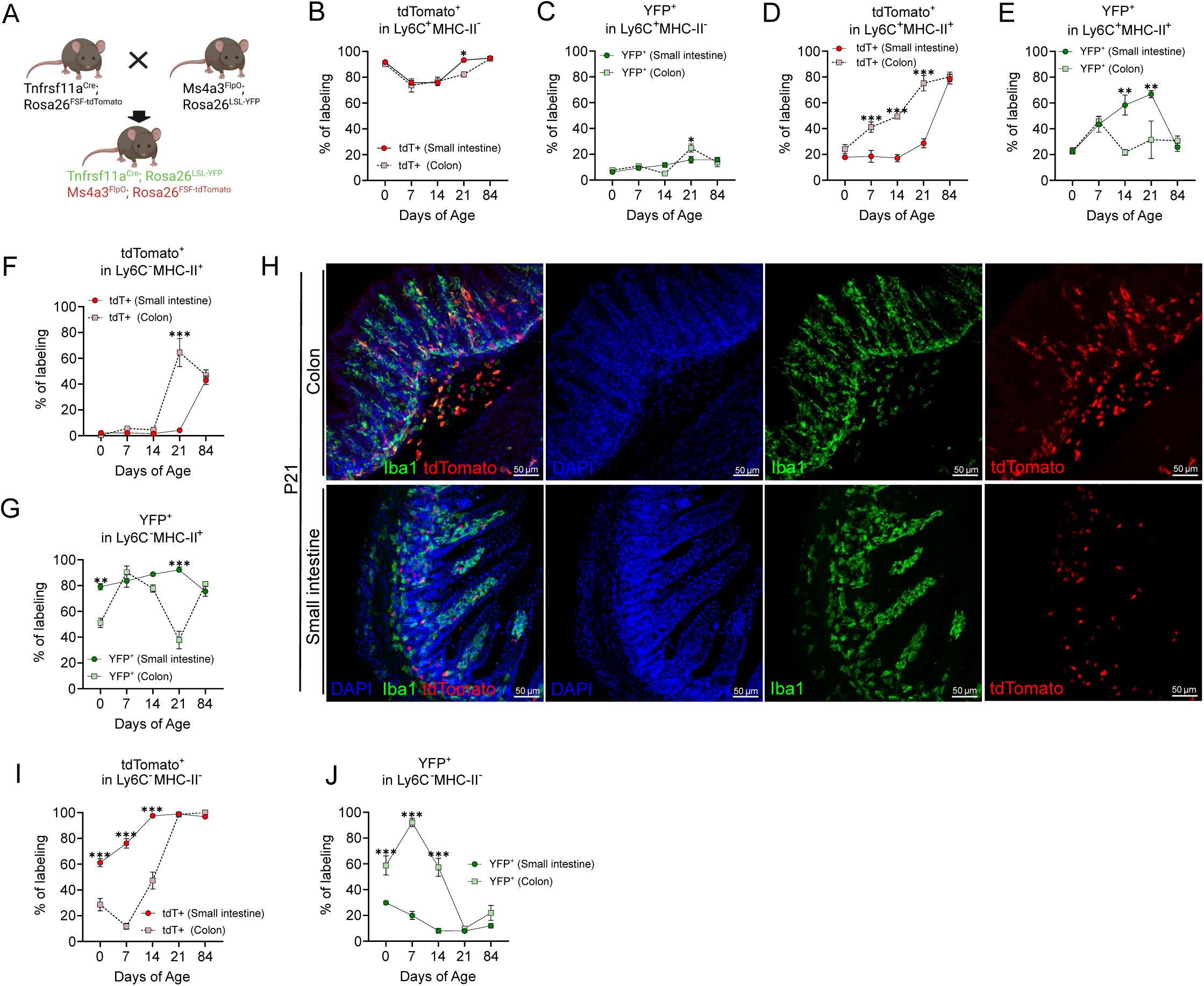
Distinct ontogenic dynamics of tissue-resident macrophages in small intestine and colon. **A.** Schematic representation of the macrophage lineage-tracing strategy using the double fate-mapping model *Tnfrsf11a^Cre^; Rosa26^LSL-YFP^; Ms4a3^FlpO^; Rosa26^FSF-tdTomato^*. *Tnfrsf11a is expressed in pre-macrophages, therefore labelling yolk sac derived macrophages YFP^+^. Ms4a3* is expressed in granulocyte–monocyte progenitors (GMPs) derived from hematopoietic stem cells (HSCs) during definitive hematopoiesis. Consequently, tdTomato⁺ cells represent short-lived or recently differentiated monocyte-derived macrophages. Long-lived monocyte-derived macrophages upregulate *Tnfrsf11a*, resulting in recombination of the *Rosa26^LSL-YFP^* reporter locus. **B.** Quantification of tdTomato⁺ signal by flow cytometry in Ly6C⁺ MHC-II**⁻** macrophages isolated from colon and small intestine from the P0 (n=3), P7 (n=5), P14 (n=3), P21 (n=4), P84 (n=3). **C.** Flow cytometric quantification of YFP⁺ cells, within the Ly6C⁺ MHC-II⁻ macrophage population isolated from colon and small intestine at P0 (n=3), P7 (n=5), P14 (n=3), P21 (n=4), P84 (n=3). **D.** Quantification of tdTomato⁺ signal, by flow cytometry in Ly6C⁺ MHC-II^+^ macrophages isolated from colon and small intestine from the P0 (n=3), P7 (n=5), P14 (n=3), P21 (n=4), P84 (n=3). **E.** Flow cytometric quantification of YFP⁺ cells within the Ly6C⁺ MHC-II^+^ macrophage population isolated from colon and small intestine at P0 (n=3), P7 (n=5), P14 (n=3), P21 (n=4), P84 (n=3). **F.** Quantification of tdTomato⁺ signal by flow cytometry in Ly6C^-^ MHC-II^+^ macrophages isolated from colon and small intestine from the P0 (n=3), P7 (n=5), P14 (n=3), P21 (n=4), P84 (n=3). **G.** Flow cytometric quantification of YFP⁺ cellswithin the Ly6C^-^ MHC-II^+^ macrophage population isolated from colon and small intestine at P0 (n=3), P7 (n=5), P14 (n=3), P21 (n=4), P84 (n=3). **H.** Immunofluorescence staining of the fate-mapping model Tnfrsf11a^Cre^; Rosa26^LSL-YFP^; Ms4a3^FlpO^; Rosa26^FSF-tdTomato^ mice, Iba1(green), Ms4a3 (red), and DAPI (blue) in the colon and small intestine of P21. Scale bar: 50 µm. **I.** Quantification of tdTomato⁺ signal by flow cytometry in Ly6C^-^ MHC-II^-^ macrophages isolated from colon and small intestine from the P0 (n=3), P7 (n=2), P14 (n=3), P21 (n=4), P84 (n=3). **J.** Flow cytometric quantification of YFP⁺ cells within the Ly6C^-^ MHC-II^-^ macrophage population isolated from colon and small intestine at P0 (n=3), P7 (n=5), P14 (n=3), P21 (n=4), P84 (n=3).

### Impact of gut microbiota on intestinal macrophage composition

We next sought to determine how the gut microbiota influences the composition of macrophage populations in the colon and small intestine. To investigate this, we analyzed macrophages in germ-free (GF) and specific pathogen-free (SPF) mice at P21. In the colon, there was no significant difference in the total number of macrophages between GF and SPF mice **(Figure 4A)**. However, the relative proportions of macrophage subsets differed. Specifically, GF mice exhibited reduced numbers of Ly6C^+^ MHC-II^-^ monocytes and Ly6C^+^ MHC-II^+^ macrophages, which consequently resulted in an increased proportion of mature resident macrophages **(Figure 4A)**. In contrast, in the small intestine, the total number of tissue-resident macrophages was significantly reduced in GF mice compared to SPF controls **(Figure 4B)**. Moreover, analysis of macrophage subsets also revealed an increased proportion of MHC-II⁺ macrophages in GF mice at 3 weeks of age **(Figure 4B)**. Similar trends were observed in 6-week-old adult mice, with an even more pronounced reduction in macrophage numbers in the small intestine of GF mice **(Figure 4C, D)**. Immunofluorescence staining confirmed a substantial decrease in MHC-II^+^ F4/80^+^ macrophages in the small intestine, but not the colon, of GF mice **(Figure 4E, F)**. Overall, our findings revealed that the microbiota had a region-specific impact on gut macrophage homeostasis. Microbial cues appeared to play a critical role in maintaining the overall macrophage pool in the small intestine, likely through modulation of monocyte recruitment and differentiation, or local macrophage renewal.

**Figure 4.**
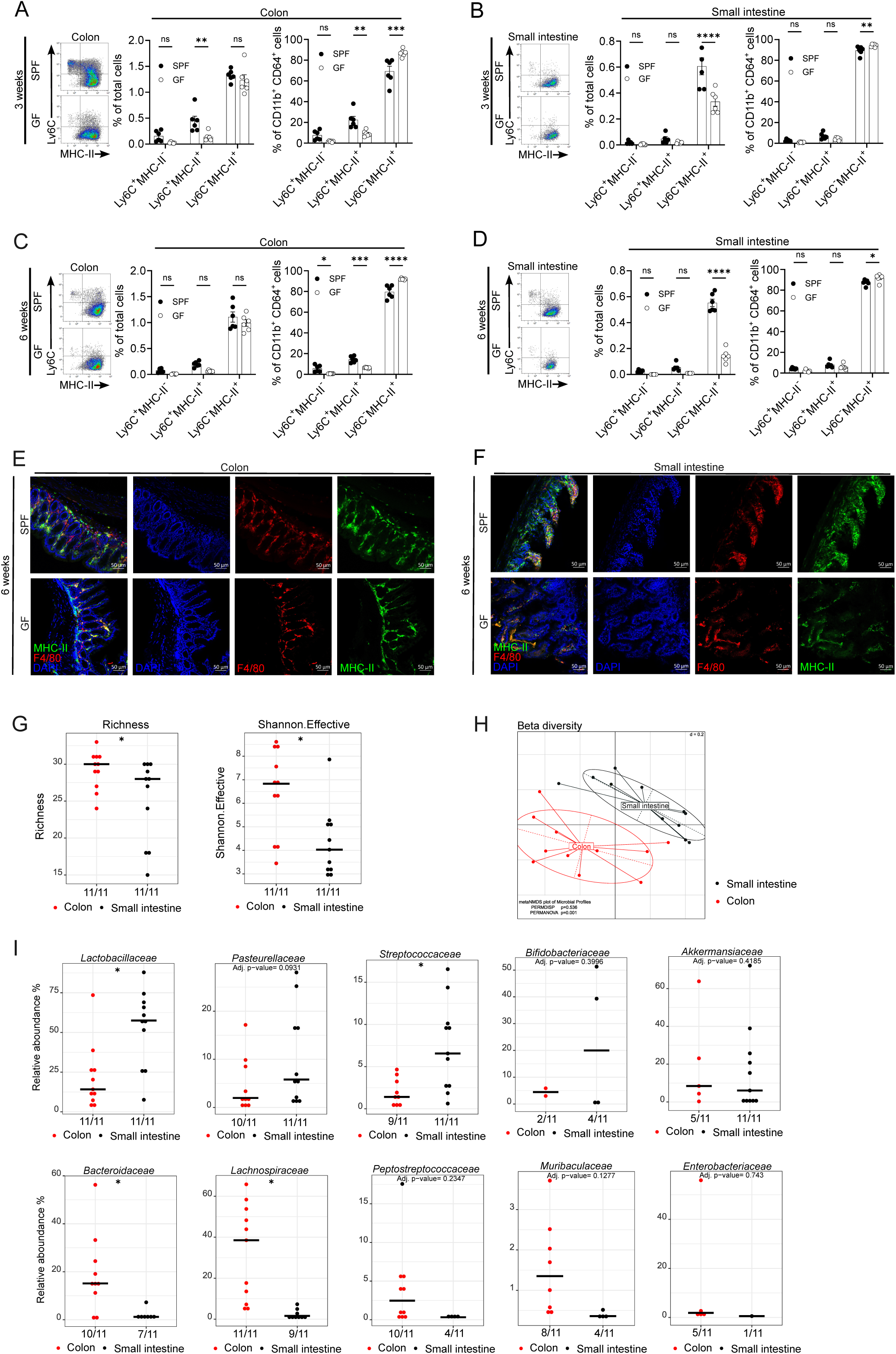
Impact of gut microbiota on intestinal macrophage composition. **A.** Quantification of macrophage populations in the colon of 3-week-old WT mice raised under specific pathogen-free (SPF) or germ-free (GF) conditions by flow cytometry. Data represent (n=6) mice per group, pooled from 2 independent experiments. **B.** Quantification of macrophage populations in the small intestine of 3-week-old WT mice raised under SPF or GF conditions by flow cytometry. Data represent (n = 6) mice per group, pooled from 2 independent experiments. **C.** Quantification of macrophage populations in the colon of 6-week-old WT mice raised under SPF or GF conditions by flow cytometry. Data represent (n =6) mice per group, pooled from 2 independent experiments. **D.** Quantification of macrophage populations in the small intestine of 6-week-old WT mice raised under SPF or GF conditions by flow cytometry. Data represent (n = 6) mice per group, pooled from 2 independent experiments. **E.** Immunofluorescence staining of MHC-II (green), F4/80 (red), and DAPI (blue) in the colon of 6 weeks old WT, SPF, and GF mice. Scale bar: 50 µm **F.** Immunofluorescence staining of MHC-II (green), F4/80 (red), and DAPI (blue) in the small intestine of 6 weeks old WT, SPF, and GF mice. Scale bar: 50 µm **G.** *Alpha*-diversity analysis. Dot plots of richness and Shannon effective counts between mouse gut regions. Statistics: Stars indicate statistical differences using the Wilcoxon signed-rank (paired) test ∗, adjusted P value (Benjamini-Hochberg (1995) correction) <0.05. **H.** *Beta*-diversity analysis. Non-metric MultiDimensional Scaling (NMDS) plot based on generalized UniFrac distances. **I.** Dot plots of microbiota composition at the family level. Differences in relative abundances (%), were assessed using a Wilcoxon signed-rank (paired) test. Numbers below the x-axis indicate the number of positive samples out of the total samples (positive/total). Adjusted p-values <0.05 were considered statistically significant. * adjusted p-value (Benjamini-Hochberg (1995) correction) <0.05. Statistical analyses and the generation of original graphs were performed in the R programming environment using Rhea ^73^. The relative abundance cut-off was set at 0.25%, and the prevalence cut-off at 30%. Abbreviations: OTU: operational taxonomic unit; NMDS: non-metric multidimensional scaling.

We hypothesized that these changes were linked to differences in microbiota profiles along the gut during development. Whilst the colon and small intestine are known to harbor distinct microbial communities in adult ^31,32^, knowledge in early life is limited. Hence, we performed 16S rRNA gene amplicon sequencing of luminal contents from the neonatal small intestine and colon. At P12, richness and Shannon effective counts were generally lower in the small intestine **(Figure 4G)**. Beta-diversity analysis showed that microbiota profiles in the colon and small intestine were distinct **(Figure 4H)**. At the family level, the relative abundance of *Lactobacillaceae*, *Pasteurellaceae*, and *Streptococcaceae* was higher in the small intestine, whereas *Bacteroidaceae*, and Lachnospiraceae were lower **(Figure 4I)**. Accordingly, the overall differences in microbial composition between colon and small intestine were already substantial at P12. In combination with other region-specific factors, these differences may contribute to the pronounced differences in macrophage populations observed between the compartments.

### Regional-specific alteration of the intestinal macrophages compartment in response to GBS

To delineate the tissue-specific immune responses to GBS in more depth, we profiled the immune cell composition in the small intestine and colon of WT mice at postnatal day 14 (P14)-two days after colonization at P12 **(Figure 5A)**. In the gut, Ly6C⁺MHC-II⁻ monocytes continuously infiltrate the LP, where they differentiate into tissue-resident macrophages ^13^. The colonic monocyte/macrophage compartment remained largely unaltered during GBS colonization **(Figure 5B)**, whereas the small intestine showed a significant reduction in Ly6C^-^macrophages upon GBS colonization **(Figure 5C)**. In contrast, GBS colonization did not alter neutrophil or eosinophil frequencies in the colon and small intestine at two days post-colonization **(Figure S2A, B, and C).** Consistently, histological analysis did not reveal infiltration of CD3ε⁺ T cells, CAE⁺ granulocytes, or B220⁺ B cells in either organ **(Figure S2D)**. At 4 days after colonization, we still observed a reduction of macrophage in the small intestine, whereas macrophage frequencies in the colon remained unchanged **(Figure S2E, F, and G)**. Moreover, granulocyte numbers were not affected by GBS colonization in either the colon or the small intestine at this time point **(Figure S2H, I)**.

**Figure 5.**
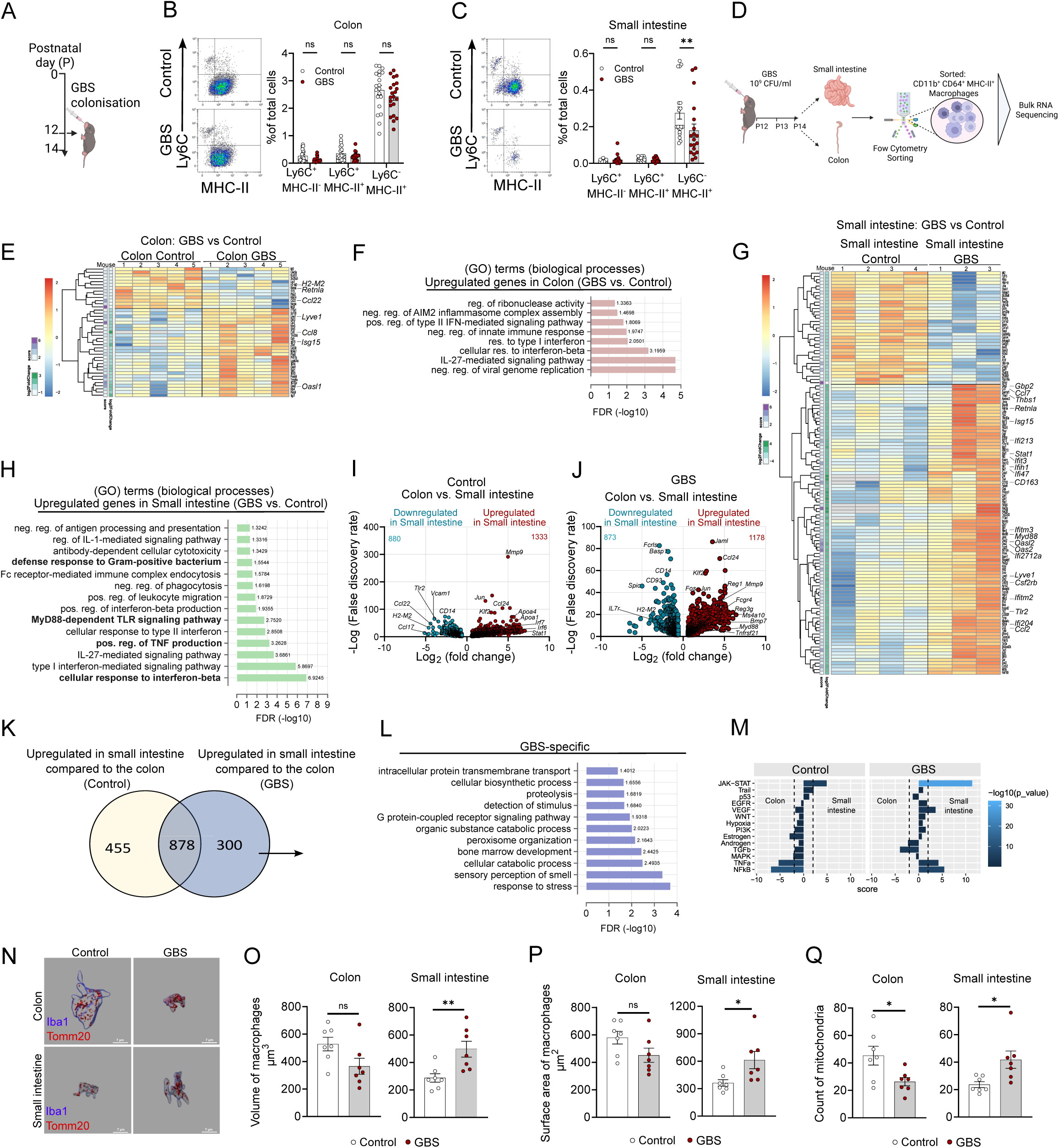
Regional-specific alteration of the intestinal macrophages compartment in response to GBS. **A.** Experimental plan for colonization and analysis at 2 days post-colonization. **B.** The frequency of Ly6^Chigh^ monocytes and MHC-II^+^ colonic macrophages among CD11b^+^ CD64^+^ cells in the colon of WT mice, 2 days post-colonization at P14, was assessed in control (n=20) and GBS-colonized mice (n=21) across 4 individual experiments. **C.** The frequency of Ly6C^high^ monocytes and MHC-II^+^ small intestinal macrophages among CD11b^+^ CD64^+^ cells in the small intestine of WT mice, 2 days post-colonization at P14, was assessed in control (n=20) and GBS-colonized mice (n=19) across 3 individual experiments. **D.** Experimental plan for bulk RNA-seq of intestinal CD11b⁺ Ly6G⁻ SiglecF⁻ CD64⁺ Ly6C⁻ MHC-II⁺ macrophages at P14, 2 days post-infection (dpi), from the colon and small intestine of C57BL/6 WT mice. **E.** Heatmap showing differentially expressed genes (DEGs) identified in colonic macrophages from mice with and without GBS colonization. DEGs were defined by an absolute fold change ≥ 1.5 (|log₂FC| ≥ log₂(1.5)) and an adjusted *p*-value (padj) < 0.05. Colonic macrophage samples were obtained from 5 GBS-colonized mice and 5 uncolonized (control) mice. **F.** GO enrichment analysis of biological process (BP) terms for differentially expressed genes (DEGs) that were upregulated in the colon of GBS-colonized samples compared to control samples. using PANTHER (https://pantherdb.org/). **G.** Heatmap depicting differentially expressed genes (DEGs) in small intestinal macrophages from mice with or without GBS colonization. DEGs were identified using an absolute fold change ≥ 1.5 (|log₂FC| ≥ log₂(1.5)) and an adjusted p-value (padj) < 0.05. Samples were initially collected from 3 GBS-colonized mice and 5 uncolonized control mice; however, 2 samples from GBS-colonized mice and 1 control sample were excluded due to insufficient RNA quality, leaving high-quality RNA from 1 GBS-colonized mouse and 4 control mice for analysis. **H.** GO enrichment analysis of biological process (BP) terms for differentially expressed genes (DEGs) that were upregulated in the small intestine of GBS-colonized samples compared to control samples, using PANTHER (https://pantherdb.org/). **I.** Volcano plot of DEGs comparing the small intestine and colon under control conditions: 1333 genes upregulated and 880 genes downregulated in the small intestine compared to the colon. **J.** Volcano plot of DEGs comparing the small intestine and colon under GBS colonization: 1178 genes upregulated in the small intestine and 873 genes downregulated in the small intestine under GBS condition, compared to the colon. **K.** Venn diagram of upregulated genes in the small intestine compared to the colon under both control and GBS colonization conditions. From figures I and J: 1333 upregulated genes in the small intestine under control and 1178 under GBS condition. Venn diagram created using GeneVenn (https://www.bioinformatics.org/gvenn/). **L.** Gene ontology analysis of the 300 genes uniquely upregulated in the small intestine compared to the colon under GBS conditions, using PANTHER (https://pantherdb.org/). **M.** PROGENy barplot is based on the DEG statistics i.e. weighted by the *stat* variable, which is the statistics variable for *padj* and *log2FoldChang.* The scoring follows the DEG direction, i.e. GBS vs <u>control</u> for groups Colon and SI and SI vs <u>Colon</u> for groups GBS and control. The underlined conditions mark the ‘baseline’, i.e. scores are enrichment for the underlined condition. **N.** Three-dimensional (3D) reconstruction of colonic and small intestinal macrophages at 2 days post–GBS colonization. Macrophages are labeled with Iba1 (blue), and mitochondria are visualized using Tomm20 (red) immunostaining. Scale bar: 7 µm. **O.** Quantification of the 3D reconstructed images using Imaris software showing the calculated macrophage volume (µm³) in the colon and small intestine at 2 days post–GBS colonization. **P.** Quantification of the 3D reconstructed images using Imaris software showing the calculated surface area of macrophages (µm2) in the colon and small intestine at 2 days post–GBS colonization. **Q.** Quantification of the 3D reconstructed images using Imaris software showing the calculated mitochondria count in the colon and small intestine at 2 days post–GBS colonization.

To investigate transcriptional alterations in intestinal macrophages upon GBS colonization, small intestinal and colonic macrophages were sorted two days after GBS colonization, and RNA from these populations was processed using the SMART-seq protocol for bulk RNA-seq analysis **(Figure 5D)**. Upon GBS colonization, colonic macrophages exhibited upregulation of only a small subset of differentially expressed genes (DEGs), including *Ccl8*, *Isg15, and Lyve1* **(Figure 5E).** Gene Ontology (GO) analysis indicated enrichment of terms such as “negative regulation of viral genome replication”, “IL27 mediated signaling pathway”, and “Type I and II interferon mediated signaling pathways” **(Figure 5F)**. In contrast, small intestinal macrophages exhibited a broader transcriptional response, characterized by the upregulation of multiple DEGs, including *Ccl2, Ccl7, Tlr2, Myd88*, and interferon family members such as *Isg15, Ifi213, Stat1, Ifit3, Ifih1, Ifi47*, *Oasl2, Oas2* **(Figure 5G)**. Some of these genes were further validated by RT-PCR, which demonstrated the same expression patterns **(Figure S2J)**. These genes were associated with GO terms like “cellular response to interferon-beta”, “positive regulation of TNF production”, “MyD88-dependent TLRs signaling pathway”, and “defense response to Gram-positive bacterium” **(Figure 5H)**. Already at baseline we observed substantial transcriptional differences between macrophages in the small and large intestine with 2,213 DEGs **(Figure 5I)**. Upon GBS colonization, 2,051 genes were up- or downregulated in the small intestine relative to the colon **(Figure 5J)**. Among all upregulated genes in the small intestine, 300 were uniquely upregulated after GBS colonization, indicating a GBS-specific transcriptional response **(Figure 5K).** GO enrichment analysis revealed significant overrepresentation of terms related to “response to stress” and “detection of stimulus” alongside “catabolic and biosynthetic processes”, suggesting that these cells are undergoing both heightened environmental sensing ^33,34^ and metabolic/functional remodeling ^35,36^. Taken together, these data support a model in which intestinal myeloid cells respond to local cues via chemo sensing and simultaneously engage in enhanced stress-adaptive metabolism (via catabolism, proteolysis, peroxisomal organization) **(Figure 5L)**. This model is consistent with an activation or stress-adapted state rather than a homeostatic one. PROGENy pathway analysis revealed that GBS colonization induced marked pathway activation, particularly in the small intestine. The small intestinal samples displayed a strong positive enrichment of the JAK–STAT pathway, representing the most prominent signaling change across all conditions. Additional activation of TNFα, MAPK, NFκB, EGFR, and VEGF pathways was also evident, indicating a broad pro-inflammatory and cytokine-driven response **(Figure 5M).** Since macrophage activation requires membrane expansion and mitochondrial biogenesis, we sought to investigate how these processes specifically influence the volume and surface area of intestinal macrophages^37^. GBS colonization induced distinct morphological alterations in small intestine macrophages, characterized by an increase in macrophage volume and surface area, as well as number mitochondria positive for Tomm20 a receptor in the mitochondrial outer membrane that recognizes and binds N-terminal targeting sequences of nuclear-encoded precursor proteins, facilitating their import into mitochondria ^38^ **(Figure 5N-Q)**. In summary, macrophages from the small intestine of neonatal mice exhibited heightened responsiveness to GBS as compared to macrophages from the colon, suggesting a compartment-specific immune regulation that may influence the course of neonatal GBS infection.

### Differential impact of MyD88 deficiency on macrophages in the small intestine

Since small intestinal macrophages exhibited a markedly stronger transcriptional response to GBS, and Myd88 was one of the significantly upregulated genes in colonization **(Figure 5G, H)**, we explored the cytological consequences of *Myd88* deficiency. Flow cytometric analysis revealed that the absence of *Myd88* markedly affected macrophage populations in the small intestine, whereas the colon was less affected. Specifically, *Myd88^⁻/⁻^* small intestines exhibited a significantly reduced total number of CD64⁺ and F4/80⁺ macrophages compared to WT controls in adult mice **(Figure 6A, and S3A)**. This reduction was predominantly observed in the Ly6C^-^ MHC-II⁺ macrophage subset in the small intestine **(Figure 6B)**, suggesting that MyD88 signaling was particularly important for maintaining this population under steady-state conditions. Immunofluorescence staining confirmed a substantial decrease in MHC-II⁺ macrophages in the small intestine, but not the colon, of *Myd88^⁻/⁻^* mice **(Figure 6C)**. Moreover, the MFIs of CD64 and MHC-II were elevated in small intestinal macrophages of *Myd88^⁻/⁻^* mice, while remaining similar between genotypes in the colon **(Figure 6D, E)**. Additionally, neutrophil and eosinophil frequencies in the colon and small intestine did not differ significantly between *Myd88^⁻/⁻^* and wild-type controls **(Figure S3B, C)**. A similar pattern was observed in P14 neonatal mice: *Myd88^⁻/⁻^* small intestines showed a marked reduction in total CD64⁺ macrophages relative to WT controls **(Figure 6F)**. This decrease was largely confined to the MHC-II⁺ subset, indicating that *Myd88^⁻/⁻^* signaling is critical for maintaining this population under steady-state conditions **(Figure 6G)**. Additionally, MFIs of MHC-II were increased in small-intestinal macrophages from *Myd88^⁻/⁻^* mice **(Figure 6H)**. In conclusion, these findings indicated that *Myd88* plays a tissue-specific role in macrophage homeostasis, exerting a dominant effect in the small intestine, but not in the colon, even before the onset of microbial colonization at birth.

**Figure 6.**
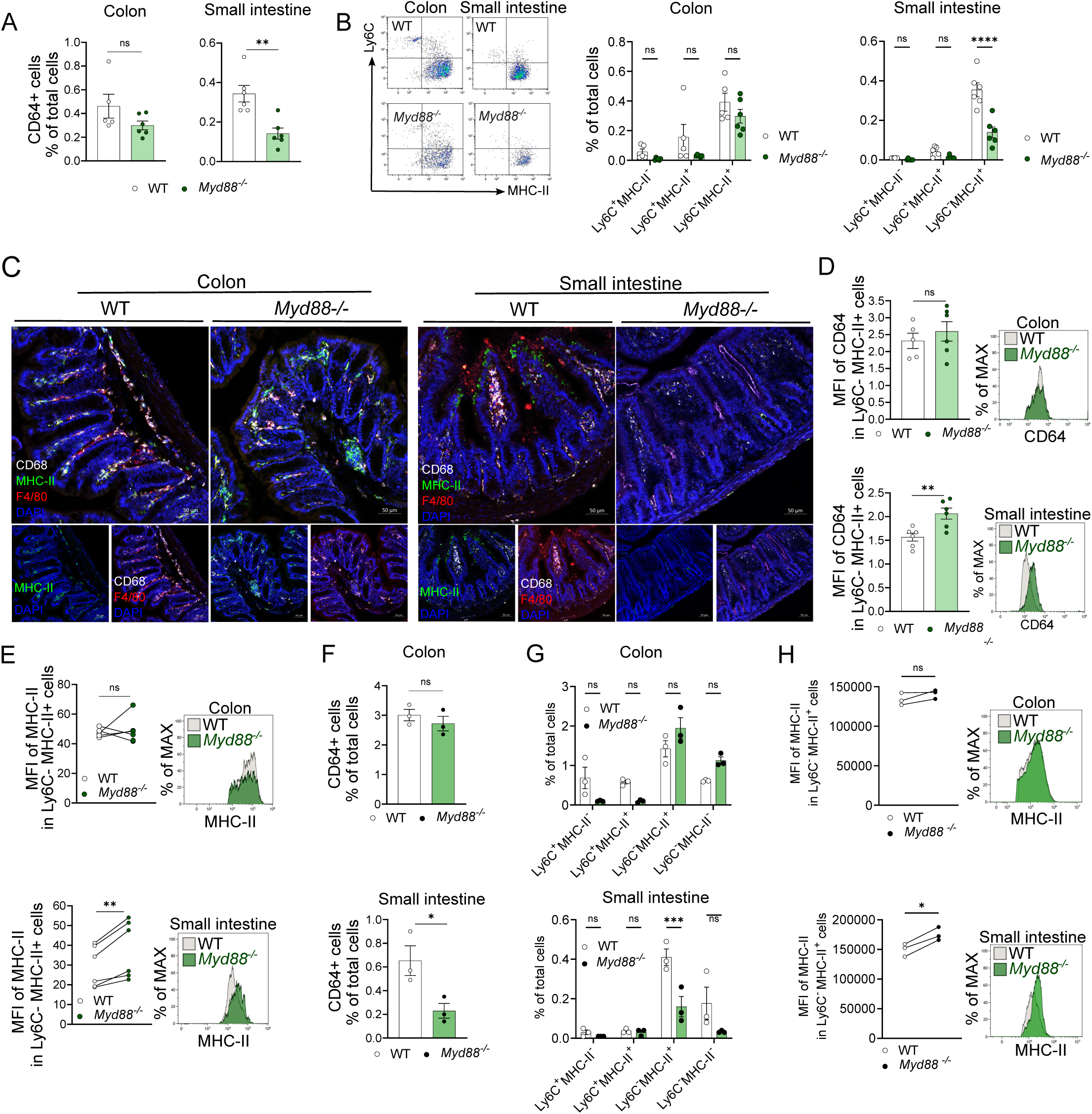
Differential impact of Myd88 deficiency on macrophage populations in the small intestine. **A.** Quantification of the CD64⁺ cells in the colon and small intestine of WT and *Myd88^⁻/⁻^* adult mice at steady state by flow cytometry. WT (n = 6) and *Myd88⁻^/⁻^* (n = 7) 6 weeks old mice were used in two independent experiments. **B.** Frequency of Ly6C⁻ MHC-II⁺ colonic and small intestinal macrophages among CD11b⁺ CD64⁺ cells in the colon and small intestine of WT and *Myd88^⁻/⁻^* mice. **C.** Immunofluorescence images of distal colon and distal small intestine, stained for DAPI (blue), MHC-II (green), F4/80 (red), and CD68 (white). Scale bar: 50 μm. **D.** Quantification of the MFI of CD64 expression in Ly6C⁻ MHC-II⁺ cells in the colon and small intestine of WT and *Myd88^⁻/⁻^* adult mice at steady state. **E.** Quantification of the MFI of MHC-II expression in Ly6C⁻ MHC-II⁺ cells in the colon and small intestine of WT and *Myd88^⁻/⁻^* adult mice at steady state. **F.** Quantification of the CD64⁺ cells in the colon and small intestine of WT and *Myd88^⁻/⁻^* adult mice at steady state by flow cytometry. WT (n = 3) and *Myd88⁻^/⁻^* (n = 3), 2 weeks old mice. **G.** Frequency of Ly6C⁻ MHC-II⁺ colonic and small intestinal macrophages among CD11b⁺ CD64⁺ cells in the colon and small intestine of WT and *Myd88^⁻/⁻^* mice. **H.** Quantification of the MFI of MHC-II expression in Ly6C⁻ MHC-II⁺ cells in the colon and small intestine of WT and *Myd88^⁻/⁻^*.

### MyD88 signaling in myeloid cells is critical for controlling GBS colonization

We next investigated whether MyD88 played a direct role in mediating protection in GBS colonization. To address this, we colonized *Myd88^-/-^*and *Myd88^+/-^* mice at P12 with GBS **(Figure 7A)**. The lethality rate was extremely high in *Myd88^-/-^* mice, with 100% succumbing to GBS within five days post-colonization **(Figure 7B)**. These mice displayed extensive GBS dissemination to the spleen, liver and brain **(Figure 7C)**, accompanied by high bacterial loads in the bloodstream **(Figure 7D)**. We then asked whether this pronounced susceptibility was restricted to neonatal mice or also observed in adults. Accordingly, we infected adult *Myd88^-/-^*mice and monitored them for three weeks post-colonization **(Figure 7E)**. Interestingly, only approximately 40% of adult *Myd88^-/-^* mice survived beyond 21 days post colonization **(Figure 7F)**, indicating that the protective role of MyD88 was not limited to early life. Moreover, the bacterial load in the fecal samples of *Myd88^-/-^* mice remained substantially higher than those in wild-type littermates’ controls **(Figure 7G)**. To determine whether inflammasome activation was the critical MyD88-mediated immune pathway involved in GBS control, we repeated the experiments with mice deficient in the adaptor protein ASC that is necessary for inflammasome activation. *Asc^-/-^* mice did not differ substantially from WT mice, indicating that inflammasome activation did not play a major role in controlling intestinal GBS colonization and translocation **(Figure S4A, B)**.

**Figure 7.**
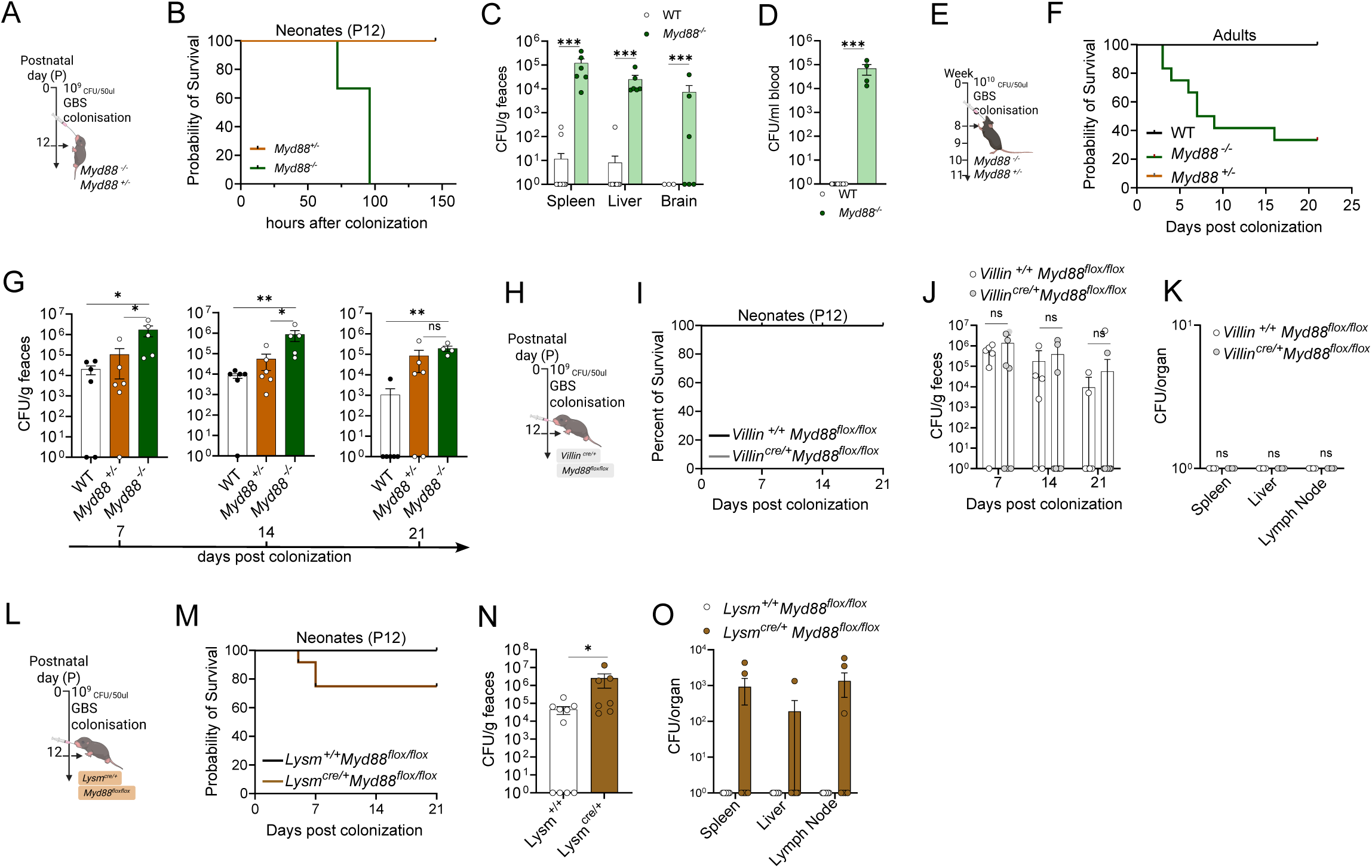
Myd88 signaling in myeloid cells is critical for controlling GBS colonization. **A.** Schematic of the experimental plan. Neonatal mice *Myd88^-/-^*, and *Myd88^+/-^* were infected at P12 with 10^9^ colony forming unit (CFU)/ml GBS via oral gavage. **B.** Survival rate following GBS colonization at postnatal day 12 in *Myd88⁻/⁻* and *Myd88⁺/⁻* mice. *Myd88^⁻/⁻^* (n= 3), and *Myd88^⁺/⁻^* (n= 3). **C.** Dissemination of GBS into organs (spleen, liver, brain/whole brain) at 2 days post colonization in neonatal mice; Spleen and liver: WT (n= 35), and *Myd88^⁻/⁻^* (n= 6), and brain: WT (n= 3), and *Myd88^⁻/⁻^* (n= 6) **D.** Quantification of the GBS Log₁₀ CFU in the blood (per ml) at 2 days post colonization in neonatal mice, WT (n= 25), *Myd88⁻/⁻* (n=4). **E.** Schematic of the experimental plan. Adult mice *Myd88^-/-^*, and *Myd88^+/-^* were infected at P12 with 10^10^ CFU/ml GBS via oral gavage. **F.** Survival rate following GBS colonized adult mice. WT (n= 9), *Myd88⁻/⁻* (n=12), and *Myd88⁺/⁻* (n=6). **G.** Log₁₀ CFU count per gram of feces from *Myd88⁻/⁻* and *Myd88⁺/⁻*, and WT mice at 7-, 14-, or 21-days post GBS colonization. WT (n=6), *Myd88⁻/⁻* (n=4), and *Myd88⁺/⁻* (n=6). **H.** Schematic of the experimental plan. Neonatal *Villin^cre/+^ Myd88^flox/flox^* mice were infected at P12 with 10^9^ colony forming unit (CFU)/ml GBS via oral gavage. **I.** Survival rate after GBS colonization at postnatal day 12 in *Villin^cre/+^ Myd88^flox/flox^* and *Villin^+/+^ Myd88^flox/flox^* mice. **J.** Log₁₀ CFU count per gram of feces in *Villin^cre/+^ Myd88^flox/flox^* (n=8) and *Villin^+/+^Myd88^flox/flox^* (n=6) mice at 7-, 14-, or 21-days post GBS colonization. **K.** Dissemination of the Log₁₀ CFU count GBS into the spleen, liver, and mesenteric lymph nodes in *Villin^cre/+^ Myd88^flox/flox^* and *Villin^+/+^Myd88^flox/flox^*mice, 2 days post colonization at P12. **L.** Schematic of the experimental plan. Neonatal *Lysm^cre/+^Myd88^flox/flox^* mice were infected at P12 with 10^9^ colony forming unit (CFU)/ml GBS via oral gavage. **M.** Survival rate after GBS colonization at postnatal day 12 in *Lysm^cre/+^ Myd88^flox/flox^* and *Lysm^+/+^Myd88^flox/flox^*mice. **N.** Log₁₀ CFU count per gram of feces in *Lysm^cre/+^ Myd88^flox/flox^* (n=7) and *Lysm^+/+^Myd88^flox/flox^* (n=10) mice 2 days after GBS colonization at P12, in 2 individual experiments. **O.** Dissemination of the Log₁₀ CFU count GBS into the spleen, liver, and mesenteric lymph nodes in *Lysm^cre/+^ Myd88^flox/flox^* and *Lysm^+/+^Myd88^flox/flox^* mice, 2 days post colonization at P12.

Given that MyD88 is also expressed in intestinal epithelial cells, we next analyzed *Villin^Cre/+^:Myd88^flox/flox^* mice **(Figure 7H)**. Interestingly, deficiency of MyD88 in epithelial cells did not impact survival after GBS colonization **(Figure 7I)**. Moreover, bacterial load in feces, were comparable between *Villin^Cre/+^* and control mice **(Figure 7J)**, and no GBS dissemination to the liver or spleen was detected **(Figure 7K)**. We next employed *LysM^Cre/+^ Myd88^flox/flox^* mice to delete *Myd88* specifically in myeloid cells. We achieved a deletion efficiency of approximately 50% in intestinal macrophages **(Figure S4C)**. Administration of GBS at P12 **(Figure 7L)** revealed that the lack of *Myd88* in myeloid cells significantly impacted survival, as approximately 25% of the mice died within the first week post colonization **(Figure 7M)**. Furthermore, two days post-colonization, *LysM^Cre/+^* mice exhibited higher GBS loads in fecal samples compared to controls **(Figure 7N)**, which was accompanied by increased GBS dissemination to the spleen, liver and mesenteric lymph nodes **(Figure 7O)**. Collectively, these findings demonstrated that MyD88 signaling in myeloid cells is essential for controlling GBS infection and preventing early mortality upon colonization. Moreover, MyD88 signaling appears to be essential for maintaining small intestinal macrophage homeostasis, since the loss of MyD88 disproportionately affects small intestinal macrophages.

## Discussion

The postnatal development of the intestinal topology is associated with requirements resulting from changes in the nutrient supply and the maturation of the microbiota ^39^. In mice, the replacement of prenatal LP macrophages in the colon by progenitors from definitive hematopoiesis has become paradigmatic for understanding postnatal tissue immune remodeling ^11^. Overall macrophage density varies along the gut, with higher concentrations in the colon than in the small intestine ^40^. However, it remains largely unclear how longitudinal trajectories, leading from a sterile tissue resorbing amniotic fluid to a zoned organ harboring distinct communities of bacteria, are intertwined with local immune development, which ensures the safe microbiota integration of pathogens.

In general, global alterations in neonatal innate immunity have been associated with the perinatal predisposition to infections and indirectly to dysbiosis, i.e. changes in gut microbiota composition ^7^. GBS is a commensal bacterium that causes invasive infections outside the neonatal period only in immunocompromised humans. In newborns of colonized women around 1% will suffer an invasive infection, sometimes with long-term consequences, making the disease a rare, but expected incident ^41^. Neonatal intestinal colonization with GBS matters, since colonization may precede invasive infection in healthy infants ^42^. Moreover, mother to infant transmission of GBS can continue for weeks after delivery ^43^. The increased risk for GBS LOD in infants originating from multiple gestation pregnancies makes a primary role of transferred antibodies, i.e. adaptive immunity, rather unlikely ^44^. Since perinatal antibiotics appear to delay GBS colonization and to increase the risk of GBS LOD, it is tempting to speculate that the immediate postnatal period represents a delicate time when intestinal immunity and microbiome are favorably interwoven to establish GBS accommodation ^45,46^.

We observed that neonatal small intestinal macrophages displayed a markedly greater MyD88-dependent responsiveness to GBS than their colonic counterparts, which pointed at compartment-specific immune regulation. Furthermore, global as well as macrophage-specific MyD88 deficiency impacted both GBS colonization and invasion without substantially influencing other myeloid cell lineages. Thus, a particular contribution of small intestinal macrophages sensing GBS presence in establishing the “streptococcal niche” at the beginning of life seems intuitive. Notably, human deficiency in TLR signaling has been directly linked to GBS LOD ^47^. Moreover, the differential responsiveness may not only shape the course of neonatal GBS infection but could also have implications for other small-intestine–predominant neonatal pathologies, such as necrotizing enterocolitis (NEC), which involves the small intestine and mechanistically links TLR activation to intestinal microcirculatory perfusion ^48^.

Notably, the intestinal zonation in LP macrophage turnover and activation, both in steady state and under mucosal challenge with GBS, was paralleled by topical differences in microbiome composition well before weaning, i.e. when the nutritional complexity was low. In particular *Lactobacillaceae*, *Pasteurellaceae*, and *Streptococcaceae* were more abundant in the small intestine, whereas *Bacteroidaceae*, and *Lachnospiraceae* were relatively more frequent in the colon already at this early age. Location of GBS colonization in the lactating mother and the neonate may be important for interindividual transmission dynamics. GBS may be subject to the enteromammary pathway, whereby bacteria translocate across the maternal intestinal mucosa and are delivered by dendritic cells to the lactating mammary gland as one source of the bacteria in pre-colostrum and milk ^49^. GBS has been linked to colonization of breast milk ducts already at its initial identification, since the species name agalactiae is derived from the “lack of milk” of cows with GBS mastitis. In humans, GBS is enriched in breast milk, and breast milk likely serves as vector for the pathogen in sepsis, most notably in parallel cases in twins^50^. On the other hand, GBS in the breast milk may be indicative of heavy reciprocal GBS exchange between mother and infant ^45^ and reciprocal oral transmission from the infant to the mammary gland may promote local pathogen load. In the context of this work, it seems important that weaning, i.e. at the transition from milk to solid food uptake, that drives the emergence of a mature macrophage population in the colon, does not substantially impact on cellularity in the SI ^40^. In contrast, the turnover of fetal macrophages by progenitors from definitive hematopoiesis in the SI that is associated with both higher basal activation and increased response to GBS exposure, is induced and maintained by the acquisition of the microbiota already at birth. Notably, a distinct weaning reaction in both colon and SI that involves RORγt regulatory T cells, short-chain fatty, acids and retinoic acid has been previously discovered by Al Nabhani et al ^51^. The regulatory mechanisms between macrophages and regulatory T cells in trans remain an intriguing future subject of study.

In summary, our work substantially expands the model integrating the – in many infants - inevitable contact with GBS around birth with the postnatal development of zoned intestinal innate immunity. Macrophage density, renewal and MyD88-dependent microbial sensing in the small intestine are primed to control GBS colonization and prevent systemic spread from the first day on. Moreover, it appears that monocyte-derived macrophages carry a particularly important function in maintaining mucosal antimicrobial immunity. Overall, the discovery of the specific role of SI macrophages to mucosal immune control of GBS early in life may help to identify new targets to prevent or attenuate late-onset GBS sepsis.

## Material and methods

### Mice

We employed mouse strains on C57BL/6J and 6N background. Wild type mice (WT) were purchased from the Jackson Laboratory (USA) or Charles River (Germany). *LysM^Cre/+^ Myd88^flox/flox^* and *Villin^Cre/+^ Myd88^flox/flox^* and *Myd88^-/-^*, and *Ccr2^-/-^* mice were obtained from Jackson Laboratories. To generate the double fate-mapper ^27^*Tnfrsf11a^Cre^*^52^; *Rosa26^LSL-YFP^* (JAX stock #006148^53^); *Ms4a3^FlpO^*^27^; *Rosa26^FSF-tdTomato^* (JAX stock #032864^54^), we bred Tnfrsf11a*^Cre/+^*; Rosa26*^FSF-tdTomato/FSF-tdTomato^* animals with Ms4a3*^FlpO^*; Rosa26*^LSL-YFP/LSL-YFP^* animals. Germ free mice were obtained from the CMF in Bern Switzerland. The SPF housed The SPF housed mice were bred under specific pathogen-free conditions at the CEMT animal facilities (University of Freiburg), housed in groups of up to five mice, and maintained in 12-hour light/dark cycles with ad libitum access to food and water. Mice were sex-matched, and littermates were randomly assigned to experimental groups.

### Bacterial cultures

Prior to infection, the GBS strain BM110 was streaked onto a blood agar plate (Columbia agar supplemented with 5% sheep blood; BioMérieux) and incubated at 37°C overnight. A single colony was then inoculated into 5 ml of Todd–Hewitt broth and incubated at 37 °C overnight. The overnight culture was subsequently diluted into 200 ml of pre-warmed Todd–Hewitt broth and grown to mid-log phase. Bacterial density was estimated using an optical spectrophotometer (OD₆₀₀ = 0.4) and confirmed by serial dilution plating on blood agar. Bacteria were then washed twice with PBS and resuspended in PBS at 10⁹ CFU/ml. For neonatal mice, 50 µl of the bacterial suspension (5 × 10⁷ CFU) or vehicle control (PBS) was administered by oral gavage. For adult mice, 50 µl containing 10¹⁰ CFU or vehicle (PBS) was administered by oral gavage.

### CFU assay

Mice were euthanized by CO₂ inhalation. The liver, spleen, and brain were collected into sterile plastic bags and mechanically dissociated. To verify assay sterility, tissues from control animals were processed and plated in parallel; no bacterial growth was detected in any of the organs analyzed.

### Lamina propria (LP) cell isolation

Cells were isolated as described previously [22]. Briefly, the colon was removed cleaned and placed in ice-cold PBS. Subsequently, the tissues were opened longitudinally, cut into 1 cm long pieces, and incubated for 30 minutes in 10 mL dissociation buffer (HBSS without MgCl_2_ (Gibco) with 2 mM EDTA (Sigma)+ 10 mM HEPES(Sigma)) at 37°C with gentle shaking (180 rpm). Following incubation, the tissues were vortexed, washed twice, and incubated in a digestion buffer (HBSS without Mg/Cl_2_, 2% FCS (Gibco, catalog number 26140079), 0.3 mg/ml Collagenase Type IV (Worthington Biochemical Corporation, catalog number LS004188), 5 U/ml dispase (Corning, catalog number 354235) and 0.5 mg/ml DNAse I (Roche/Sigma, catalog number SLBV1446)) twice for 25 minutes at 37 °C and 180 Supernatant containing cells was collected through a 70µm cell strainer, which was washed between steps, in a 50 mL Falcon tube. Finally, the samples were centrifuged at 4 °C and 400g for 10 minutes. See Flow cytometric analysis and cell sorting gating strategy **(Figure S1A)**. For *Tnfrsf11a^Cre^; Rosa26^LSL-YFP^; Ms4a3^FlpO^; Rosa26^FSF-tdTomato^* lamina propria the tissue was cleaned, separated from the *muscularis externa,* and cut into 1cm long pieces. The tissue was opened longitudinally and first incubated in HBSS without MgCl_2_ (PAN Biotech) containing 1% FCS (PAN Biotech), 100 μg/mL penicillin/ streptomycin (PAN Biotech), 1 mM EDTA (Sigma), and 1 mM DTT (Sigma) for 20 minutes at 37°C with gentle shaking (220rpm). Next, tissue was washed and incubated in HBSS without MgCl_2_ (PAN Biotech) containing 1% FCS (PAN Biotech), 100 μg/mL penicillin/ streptomycin (PAN Biotech), 1 mM EDTA (Sigma) for 20 minutes at 37°C with gentle shaking (220rpm) to remove all epithelial cells. Next, tissues were washed and minced in digestion buffer containing RPMI 1640 (PAN Biotech), 5% FCS (PAN Biotech), 100 μg/mL penicillin/ streptomycin (PAN Biotech), 2% HEPES (Gibco), 100U/ml DNAse I (Sigma Cat. DN25), 1mg/ml Dispase II (Gibco, Cat. 17105041) and 100U/ml Collagenase D (Roche, Cat. 11088858001) for 20 minutes at 37°C with gentle shaking (220rpm). After digestion, cell suspensions were filtered through a 70µm cell strainer and abundantly washed with FACS buffer, followed by centrifugation at 4 °C and 400x*g* for 10 minutes.

### Flow cytometric analysis and cell sorting

To block FcγII/III receptors, cells were incubated with anti-CD16/32 antibody (eBioscience) for 5 min at 4 °C in FACS buffer (PBS, 2% FCS, 2 mM EDTA). Following Fc blockade, LP cells were stained with the appropriate antibody panel, including anti-mouse CD45 (30-F11, BioLegend), CD64 (X54-5/7.1.1, BD Biosciences), Ly6C (HK1.4, Invitrogen/eBioscience), CD11b (M1/70, eBioscience), Ly6G (1A8, BioLegend), Siglec-F (S17007L, BioLegend), MHC-II (AF6-120.1, eBioscience), and CD206 (catalog number pending). Antibody mixtures were added to the cells and incubated for 30 min at 4 °C in the dark. Cells were then washed and resuspended in 200 µl of FACS buffer for acquisition. Flow cytometry data were acquired on a Gallios™ (Beckman Coulter) or LSRFortessa™ (BD Biosciences) and analyzed using Kaluza (v2.1, Beckman Coulter) or FlowJo (v10, FlowJo LLC). Cell sorting was performed using a MoFlo Astrios AQ or a CytoFlex SRT (Beckman Coulter). Cells from *Tnfrsf11a^Cre^; Rosa26^LSL-YFP^; Ms4a3^FlpO^; Rosa26^FSF-tdTomato^* mice were first blocked with anti-CD16/32 antibody (Biolegend) for 15 min at 4 °C in FACS buffer, then incubated with an antibody cocktail containing CD45 (30-F11, BD Biosciences), CD64 (X54-5/7.1.1, Biolegend), Ly6C (HK1.4, Biolegend), CD11b (M1/70, BD Biosciences), MHC-II (M5/114.15.2, Biolegend) for 30 min at 4 °C in the dark. Cells were then washed and resuspended in 200 µl of FACS buffer and counter-stained with 7-AAD for acquisition. Flow cytometry data were acquired on a SONY ID7000 spectral analyser (SONY) and analysed using FlowJo (v10, FlowJo LLC).

### Histology & Immunofluorescence

For histopathology, distal colons, and distal small intestine were fixed in 4% paraformaldehyde (PFA) for 4 hours at 4°C, followed by immersion in 20% sucrose for at least 1 hour, or until the tissue sank. The tissue was snap-frozen in liquid nitrogen using TissueTek embedding. Sections with a thickness of 15 µm were then cut at - 18°C to −20°C for further examination. The H&E staining procedure involved washing the sample once with PBS, followed by washing with distilled water. Hematoxylin (GILL III) was applied for 4 minutes, followed by washing with tap water. A rinse with 0.1% HCl for 10 seconds was performed, followed by another wash with tap water. Eosin (1%) was applied for 30 seconds, followed by washing with tap water. The slides were immersed in 100% ethanol for 2 minutes (repeated), then in isopropanol for 2 minutes, and finally in xylene for 5 minutes (repeated). Slides were air-dried in the hood and mounted with Dako Fluorescence Mounting Media. For immunofluorescence, the washing media consisted of PBS, 1% BSA, and 0.1% Triton 10X. The protocol began with washing the sample with PBS, followed by a 40-minute incubation in the washing media. Subsequently, Hoechst (1:500), CD68 (FA-11 BioLegend), F4/80 (BM-8 eBioscience), and MHC-II (m5/114.15.2 invitrogen1) were added to the washing media and incubated for 3 hours at room temperature. For goblet cell staining, the Alcian blue staining kit (ab150662) was used. For liver histological analysis, slides were stained using Hoechst (1:500), CD68 (FA-11 BioLegend), F4/80 (BM-8 eBioscience), and iNOS (CXNFT invitrogen). Confocal microscopy was performed with a ZEISS LSM710, LSM880, or LSM980 confocal microscope equipped with a 20x/0.8 NA Plan-Apochromat objective (Carl Zeiss Microimaging) using 1-mm optical slices. Pictures were then analyzed and edited using the ZEN blue 3.0 software. For *Tnfrsf11a^Cre^; Rosa26^LSL-YFP^; Ms4a3^FlpO^; Rosa26^FSF-tdTomato^* mice distal colons, and distal small intestine were fixed in 4% paraformaldehyde (PFA) for 1 hour at 4°C, followed by immersion in 30% sucrose for at least 1 hour, or until the tissue sank. The tissue was snap-frozen in liquid nitrogen using TissueTek embedding. Sections with a thickness of 12 µm were then cut at −18°C to −20°C for further examination. Tissue was first washed 3 times in PBS, then permeabilized for 2 hours at room temperature in PBS containing 0.4% Triton-X 100. Next, tissue was blocked using PBS containing 0.4% Triton-X 100, 5% Donkey serum (PAN-Biotech), 5% BSA (Sigma) and 1% anti-CD16/32 antibody (Biolegend) (blocking buffer) for 2 hours at room temperature. Tissue was incubated overnight at 4°C with anti-Iba-1 (abcam, ab178847) in blocking buffer. After washing in PBS 3 times, tissue was then incubated for 2 hours at room temperature with donkey anti-rabbit AF555 (Invitrogen, A31573) in blocking buffer. Tissue was washed 3 times, incubated with DAPI in PBS for 10 minutes, then mounted using Fluoromount G mounting medium (Invitrogen) and imaged at a Zeiss LSM 880 confocal microscope.

### Tissue Preparation, Immunohistochemistry, and CAE Staining

Tissue samples from the small and large intestine were fixed in 4% paraformaldehyde (PFA) for 24 hours, dehydrated through a graded ethanol series, and embedded in paraffin. Sagittal sections of 3 µm thickness were prepared using a Microm HM 355S microtome and mounted on SuperFrost™ glass slides. Residual paraffin was removed by incubating the slides at 80 °C for 30 minutes. Sections were subsequently deparaffinized in xylene and rehydrated through a descending ethanol series (2× 100%, 2× 96%, 2× 70%). Antigen retrieval was carried out by incubating the slides in a pH 6.0 citrate buffer at 92 °C for 30 minutes. Following a 15-minute cooling step in a water bath, slides were transferred into phosphate-buffered saline (PBS). Endogenous peroxidase activity was quenched using 3% hydrogen peroxide (diluted 1:10 from 30% H₂O₂ stock) in PBS for 10 minutes. After several washes in Tris buffer (tris(hydroxymethyl)aminomethane-based), nonspecific binding was blocked by incubating each slide with 150 µl of blocking buffer (containing goat serum, 1% Triton X-100, and Tris buffer) for 1 hour at room temperature. Primary antibodies against CD3 (DAKO IR503) and B220 (BD Pharmingen 557390) were applied and incubated for 30 minutes at room temperature. After washing with Tris buffer, a secondary anti-rat antibody (diluted 1:200 in blocking buffer) was added for 15 minutes at room temperature. Subsequently, Streptavidin-HRP (diluted 1:1000 in Tris buffer) was applied for 20 minutes, followed by three washes with Tris buffer. Immunoreactivity was visualized using 3,3′-diaminobenzidine (DAB; 1 ml blocking buffer + 1 drop DAB). Slides were rinsed in distilled water for 5 minutes, dehydrated through an ascending ethanol series (70%, 96%, 100%), and mounted with glycerol.

Chloracetate esterase (CAE) staining was performed using the Sigma 91CKT kit (Sigma-Aldrich) following the manufacturer’s instructions.

Microscopic imaging was performed using a Keyence BZ-X800 microscope at 20× magnification.

### Tomm20 Immunostaining and 3D Image Quantification

The tissues were blocked and permeabilized for 60 minutes with 1 ml of 5% BSA + 0.1% Triton X-100 per slide in a dark box. For more precise application of the antibodies, the three sections on each slide were outlined with a fat-pen after thoroughly draining the solution. To visualize macrophages, an antibody against Iba-1 (dilution 1:250 in 5% BSA) was used. To stain mitochondria, an antibody against Tomm20 (dilution 1:100 in 5% BSA) was applied. The sections were incubated with a total volume of 50 µl at 4 °C for 48 hours in the dark. To better secure the sections, a paraffin film was placed on each slide. After the 48-hour incubation, the sections were washed three times for 10 minutes with PBS.

In the second step of the staining procedure, the Alexa488-conjugated secondary antibody (dilution 1:500 in 5% BSA) and the Alexa568-conjugated secondary antibody (dilution 1:500 in 5% BSA) were apped with a total volume of 50 µl. The sections were then incubated for another 48 hours at 4 °C in the dark. Next, the sections were washed three times for 10 minutes with PBS and subsequently stained with DAPI (dilution 1:10,000 in PBS). The slides were again washed for 10 minutes with PBS. After mounting a 25×50 mm coverslip with Mowiol, the slides were dried for another 48 hours and stored at 4 °C.

### RNA extraction and qRT-PCR

To extract RNA, cell populations were sorted in RNa lysis buffer supplemented with 1% β-mercaptoethanol. RNA extraction was carried out using either the Extractme total RNA micro spin kit or the RNeasy Micro Kit Plus (Qiagen), following the manufacturer’s instructions. Subsequently, cDNA synthesis was performed using the SuperScriptTM IV VILO mix (Thermo Fisher). For qRT-PCR, ABsolute qPCR SYBR Green from Thermo Fisher Scientific was employed, and the samples were analyzed using a LightCycler 480 (Roche).

Mouse Oligonucleotides for qRT-PCR included the following sequences (5‘-3‘):

*Gapdh* (for: ACTCCACTCACGGCAAATTC, rev: TCTCCATGGTGGTGAAGACA)

*Tnf* (for: TCGTAGCAAACCACCAAGTG, rev: CCTTGTCCCTTGAAGAGAACC)

*Cxcl1* (for: TTGCCTCAATCCTGCATCCC, rev: GTTGGATTTGTCACTGTTCAGCAT)

Intestinal macrophages were isolated from the colon and small intestine of P14 mice, 2 days after GBS colonization, and RNA was obtained using Arcturus PicoPure™ RNA Isolation Kit. Reverse transcription and real-time PCR analysis were performed using high-capacity RNA-to-cDNA-Kit and Gene Expression Master Mix reagents (Applied Biosystems) according to the manufacturer’s recommendations. RT-PCRs were analyzed with a LightCycler 480 (Roche). For gene expression analysis, the following TaqMan Gene Expression Assays were used:

*Actb* (Mm01205647_g1)

*Ccl2* (Mm00441242_m1)

*Retnla* (Mm00445109_m1)

*Tlr2* (Mm00442346_m1)

*Ccl7* (Mm00443113_m1)

### RNA-seq (SMART3-seq)

Intestinal macrophages were isolated from the colon and small intestine of P14 mice, 2 days after GBS colonization, and RNA was obtained using Arcturus PicoPure™ RNA Isolation Kit. cDNA library preparation was performed following the Smart3-Seq method described in ^55^.

### mRNA-seq analysis, DEG & Pathways

Processing of the SMART-seq bulk mRNA-seq from raw single-end fastq files to count matrix was performed using snakePipes 2.7.3^56^. In brief, reads were trimmed with standard Illumina barcode and PolyA-tails with A-stretches of 10 or longer with cutadapt^57^ and UMItools used to deduplicate the libraries using the barcode pattern NNNNNCCC. After STAR alignment ^58^ against GRCm39, featureCounts ^59^ was used to quantify gene expression from exons using--libtype 1 on Ensembl v109^60^. For the differential analysis, the DESeq2 ^61^ was used to model condition, sample sex and sequencing batch and compute log_2_ fold changes between condition. Sample sex was determined by relative Xist expression (x > 5/10000). For expression heatmaps, FPMs were computed and batch-corrected using Limma’s ^62^ by regressing out the effect of sample sex and sequencing batch. After removal of batch effects, technical replicates were averaged. PROGENy analysis^63^ was run decoupler’s univariate linear model ^64^ with the standard mouse progeny database. Genes were ranked according to the DEseq2 stat variable.

### Isolation of metagenomic DNA

The DNA from mouse faecal samples was isolated using a modified version of the protocol by ^65^. Samples were melted and homogenised in 600 µl stool DNA stabilizer (Stratec biomedical), before cell lysis was conducted by adding 500 mg of 0.1 mm-diameter silica/zirconia beads, 250 μl 4 M guanidine thiocyanate in 0.1 M Tris (pH 7.5) and 500 μl 5 % N-lauroyl sarcosine in 0.1 M PBS (pH 8.0). They were incubated at 70 °C and 700 rpm for 60 min. Finaly, the cell disruption was carried out on a FastPrep® instrument (MP Biomedicals) for 40 s at 6.5 M/s, 3 times under cooling. 15 mg Polyvinylpyrrolidone (PVPP) were added and mixed with the samples before being centrifuged at 15.000 x g at 4 °C for 3 min. 650 µl of the supernatant were transferred into a new 2 ml tube and centrifuged for 3 min at 15.000 x g and 4 °C, before 500 µl were transferred into a new 2 ml tube containing 50 µg of RNase. Samples were incubated for 20 minutes at 37 °C and 700 rpm. gDNA was isolated using the NucleoSpin® gDNA Clean-up Kit from Macherey-Nagel according to the manufacturer’s protocol. DNA was eluted from columns twice using 40 µl Elution buffer and concentration was measured with NanoDrop® (Thermo Scientific). Samples were stored at - 20 °C. A blank extraction have been used as a negative control.

### 16S rRNA gene amplicon sequencing

Library preparation and sequencing were performed using an automation platform (Biomek400, Beckman Coulter) as described in detail previously by Lagkouvardos et al., 2015. Succinclty the V3-V4 region of 16S rRNA genes was amplified in duplicates (30 cycles) using primers 341F-785R ^66^ following a two-step protocol and dual indexing ^67^. The AMPure XP system (Beckman Coulter) was used for purification. Sequencing of pooled samples in paired-end modus (PE300) was carried out on a MiSeq system (Illumina, Inc.) with 25% (v/v) PhiX standard library according to the manufacturer’s instructions.

### 16S rRNA gene amplicon data analysis

Processing of raw reads at the level of molecular strains (ZOTUs) was done using an updated version of an in-house developed pipeline (www.imngs.org) ^68^ using the UNOISE3 method ^69^ from USEARCH11 ^70^. Sequences were demultiplexed and trimmed to the first base with a quality score >20. Sequences with less than 350 (200 for V4 only) and more than 500 nucleotides and paired reads with an expected error >3 (>2 for V4 only) were excluded from the analysis. Remaining reads were trimmed by 20 nucleotides (10 nucleotides for V4 only) on each end to avoid GC bias and non-random base composition. The minimum ASV (amplicon sequence variant) size was four. Alignment and taxonomic classification of denoised sequences was conducted with SINA 1.6.1 ^71^, using the taxonomy of SILVA release 138.1 ^72^. For analysis at the level of operational taxonomic units (SOTUs), the denoised sequences (ZOTUs) from the previous step were clustered at 97% sequence similarity using USEARCH11 and only those with a relative abundance >0.25% in at least one sample were kept. Statistical analysis was performed in R using the Rhea pipeline (v1.1.6) (https://lagkouvardos.github.io/Rhea/) ^73^. To account for differences in sequence depth OTU tables were normalized. Diversity within samples (α-diversity) was assessed on the basis of species richness and Shannon effective diversity ^74^. Diversity between samples (β-diversity) was computed based on generalized UniFrac distances ^75^. Zero values were removed from statistical calculations and p-values were calculated either using a non-parametric Wilcoxon Rank Sum Test or a Fisher’s exact test corrected for multiple comparisons according to the Benjamini-Hochberg method. Only taxa with a prevalence of at least 30 % samples in one given group were considered for statistical analysis.

## Data availability

Raw data of the 16S rRNA gene amplicon sequencing were submitted to ENA and are available under accession number PRJEB103989. Bulk RNA-seq data were submitted to GEO under the accession numbers GSE314376.

## Statistical analysis

Sample size (*n*) denotes biological replicates and data points are presented as mean ± standard error of the mean (SEM) of independent experiments as indicated. Statistical analysis was performed using GraphPad Prism for Windows. Differences were considered significant when *P* < 0.05.

## Acknowledgements

The authors are grateful to Anita Imm and Adriana Greco for their great technical assistance. We are indebted to the Lighthouse facility (Freiburg) for their assistance with cell sorting and confocal microscopy and thank the Center for Experimental Models and Transgenic Service (CEMT) of University Medical Center Freiburg for excellent technical support and assistance with the animal studies performed in this work. We thank Ntana Kousetzi from Aachen University Hospital for 16sRNA sequencing. We thank Yasuhiro Kobayashi for providing *Tnfrsf11a^Cre^* mice and Florent Ginhoux for providing the *Ms4a3^FlpO^* mice. Experimental schemes were created with <u>Biorender.com</u>.

This work was funded by the Hans A. Krebs Medical Scientist Program, Faculty of Medicine, University of Freiburg (Z.M.M.) and the IMM-PACT-Program for Clinician Scientists, Department of Internal Medicine II, Medical Center—University of Freiburg and Faculty of Medicine, University of Freiburg (M.F.); the German Research Foundation (DFG) under TRR359 (#491676693, T.C., J.K., P.H., D.E.), TRR167 (#259373024, D.E., P.H.), CRC1160 (#256073931, P.H.), (HE3127/9, HE3127/12, and HE3127/16, P.H.), 413517907 (M.F.) and (LA3680/9-1 and LA3680/10-1, N.L.); under Germany’s Excellence Strategy (CIBSS—EXC-2189, #390939984, P.H.); the Deutsche Forschungsgemeinschaft (DFG, German Research Foundation) under Germany’s Excellence Strategy-EXC2151-390873048 (E.M.), SFB1454 (#432325352, E.M.), FOR5547 (#503306912, E.M.), FOR5775 (#533863915, E.M.), as well as an EMBO fellowship (ATLF 873-2023, M.F.V). D. E. was supported by the Else Kröner-Fresenius-Stiftung (EKFS) and Ministry of Science, Research and Arts, Baden-Wuerttemberg under the aegis of JPND. We would like to thank the Flow Cytometry Core Facility of the Mathematical and Natural Sciences Faculty at the University of Bonn for providing support and instrumentation funded by the Deutsche Forschungsgemeinschaft (DFG, German Research Foundation) - #553871989, #341039622.

## Authorship

Contribution: Z.M.M., M.F., J.K., D.E., and P.H. conceptualized the study and were responsible for the methodology and study design; Z.M.M., M.F., J.K., and P.H. wrote the manuscript; Z.M.M. was responsible for visualization; M.F.V., M.E., C.R., L.K., M.R., R.A., N.T., R.R.E., S.B., M.B., E.M., T.C., D.E., contributed to reviewing and editing the draft; Z.M.M., M.F., M.F.V., M.E., L.K., R.M., R.R.E., and S.B. performed the investigation; C.R., Z.M.M., N.T., and T.C. performed 16S ribosomal RNA gene sequencing; M.E. performed RNA sequencing and M.R., M.E., Z.M.M., and M.B. preformed RNA-seq data analysis; E.M. provided the mice and contributed to critical revision; D.E., J.K., and P.H. supervised the study and acquired funding.

**The authors declare no competing interests.**

## Supplementary figures

**Figure S1:**
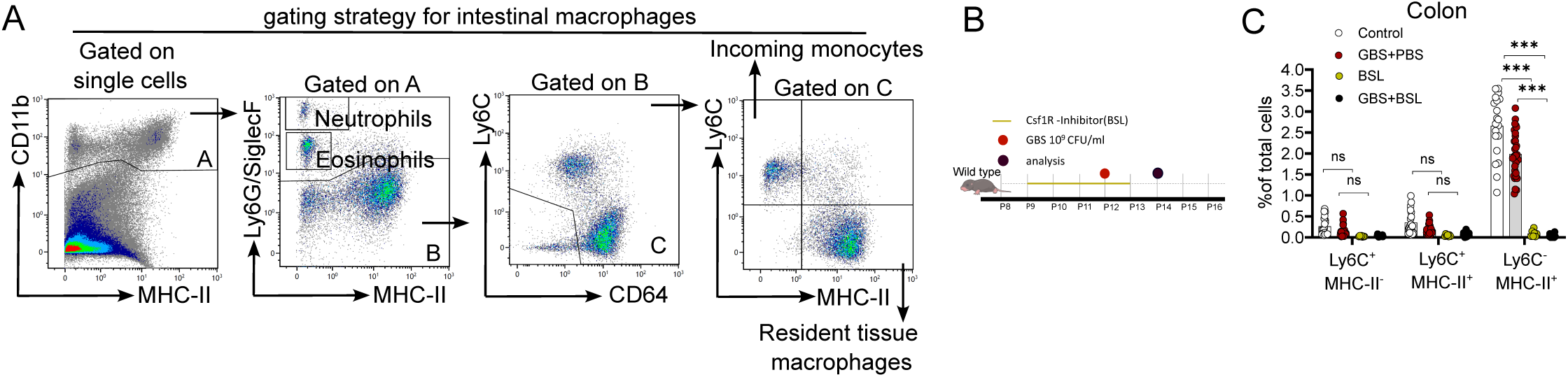
Intestinal myeloid cell analysis following CSF1 receptor inhibition. **A.** Single cells were initially gated to be CD11b+ myeloid cells (Gate A). Subsequently, Ly6G+ (neutrophils) and SiglecF+ (eosinophils), and Ly6C-CD64-cells were excluded. Cells were further subdivided into Ly6C+ infiltrating monocytes and Ly6C-MHC-II+ intestinal macrophages. **B.** Schematic representation of experimental design. WT mice received daily administration of the CSF1 receptor inhibitor (BSL) for five consecutive days, starting at postnatal day 9 (P9). Bacterial colonization was performed at P12. Experimental analyses were conducted either 2- or 4-days post-colonization. **C.** Flow cytometric assessment of macrophage depletion efficiency in the Colon. Tissue-resident macrophages were identified by sequential gating on CD45⁺ CD11b⁺ Ly6G⁻ SiglecF⁻ CD64⁺ cells, 2 days post colonization. Experimental groups included Control (n=20), GBS+PBS (n=27), BSL (n=4), and BSL+GBS (n=15), analyzed across 4 independent experiments.

**Figure S2:**
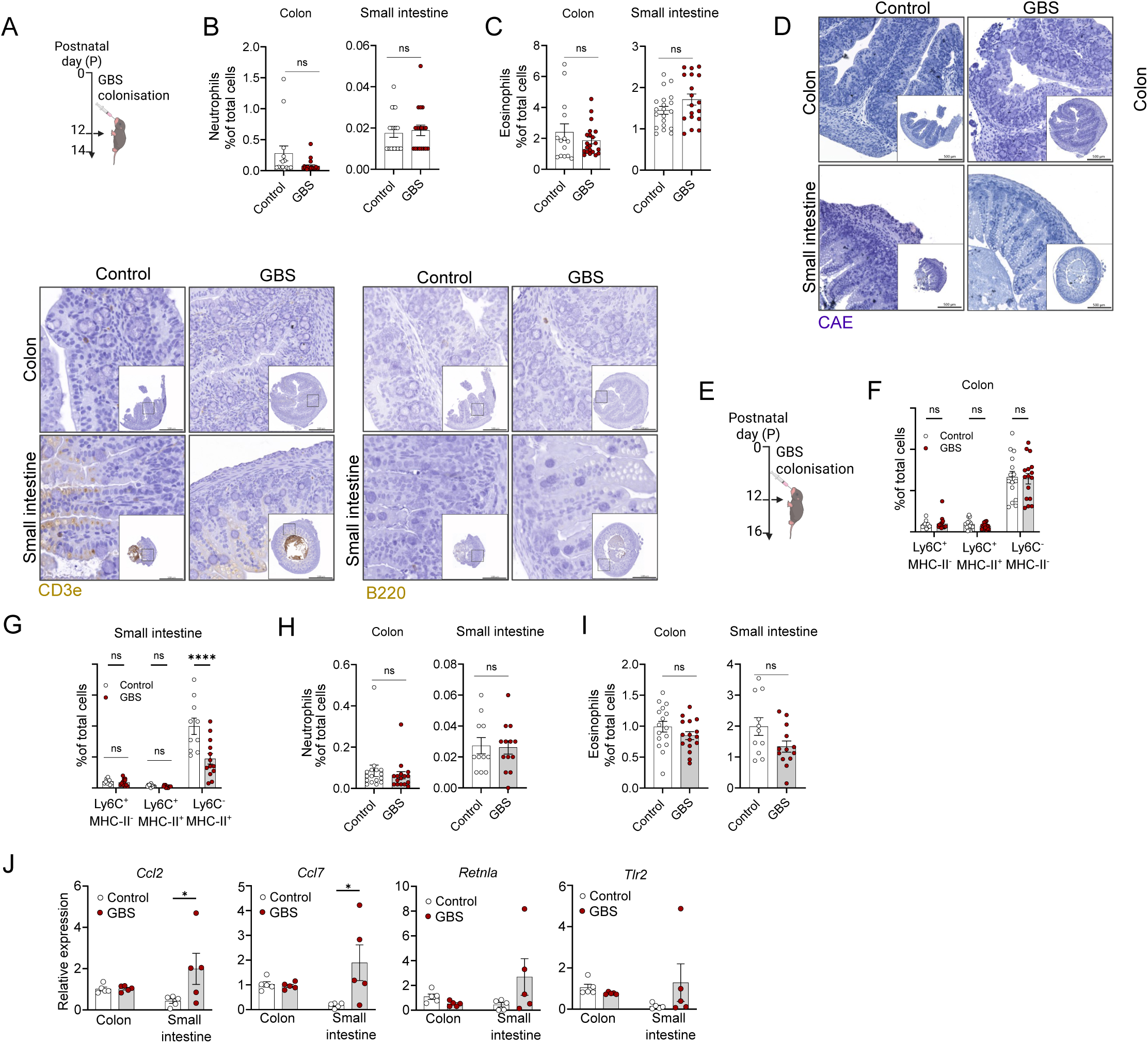
Analysis of myeloid and lymphoid immune responses following neonatal GBS colonization. **A.** Experimental plan for colonization and analysis at 2 days post-colonization at P14. **B.** Neutrophils quantification in the colon and small intestine by flow cytometric, 2 days post-colonization at P14, was assessed in control (n=14) and GBS-colonized mice (n=15) across 3 individual experiments. **C.** Eosinophils quantification in the colon and small intestine by flow cytometric, 2 days post-colonization at P14, was assessed in control (n=14) and GBS-colonized mice (n=15) across 3 individual experiments. **D.** CAE Staining, Immunohistochemistry (IHC) of CD3 and B220 in the colon and small intestine, 2 days post-colonization at P14. Scale bar indicates 500µm **E.** Experimental plan for colonization and analysis 4 days post-colonization at P16. **F.** The frequency of Ly6C^high^ monocytes and MHC-II^+^ colonic macrophages among CD11b^+^ CD64^+^ cells in the colon of WT mice, 4 days post-colonization at P16, was assessed in control (n=11) and GBS-colonized mice (n=13) across 3 individual experiments. **G.** The frequency of Ly6^Chigh^ monocytes and MHC-II^+^ small intestinal macrophages among CD11b^+^ CD64^+^ cells in the colon of WT mice, 4 days post-colonization at P16, was assessed in control (n=11) and GBS-colonized mice (n=13) across 3 individual experiments. **H.** Neutrophils quantification in the colon and small intestine by flow cytometric, 4 days post-colonization at P16. **I.** Eosinophils quantification in the colon and small intestine by flow cytometric, 4 days post-colonization at P16. **J.** Quantitative real-time PCR (qRT-PCR) analysis of *Ccl2, Ccl7, Retnla, Tlr2* gene expression in Ly6C^-^ MHC-II⁺ cells isolated from the colon (n=5) and small intestine (n=5) of the control mice and colon (n=5) and small intestine (n=5) of the 2 days post GBS colonized mice.

**Figure S3:**
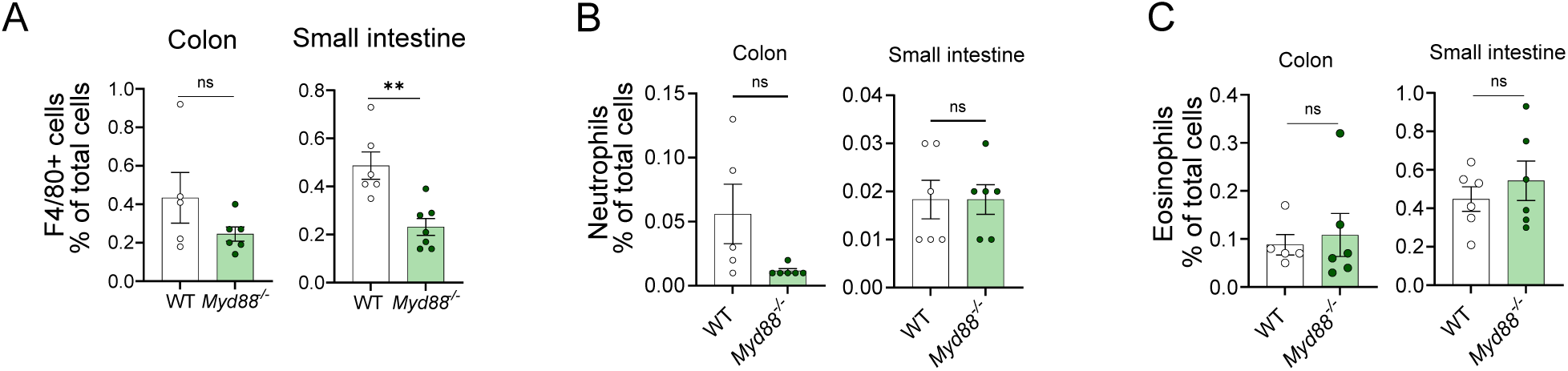
Steady-state myeloid cell populations in the colon and small intestine of WT and *Myd88^⁻/⁻^* adult mice. **A.** Quantification of the F4/80⁺ cells in the colon and small intestine of WT and *Myd88^⁻/⁻^* adult mice at steady state by flow cytometry. WT (n= 6) and *Myd88^⁻/⁻^* (n= 7) mice were used in two independent experiments. **B.** Quantification of the neutrophils in the colon and small intestine of WT and *Myd88^⁻/⁻^* adult mice at steady state by flow cytometry. **C.** Quantification of the eosinophils in the colon and small intestine of WT and *Myd88^⁻/⁻^* adult mice at steady state by flow cytometry.

**Figure S4:**
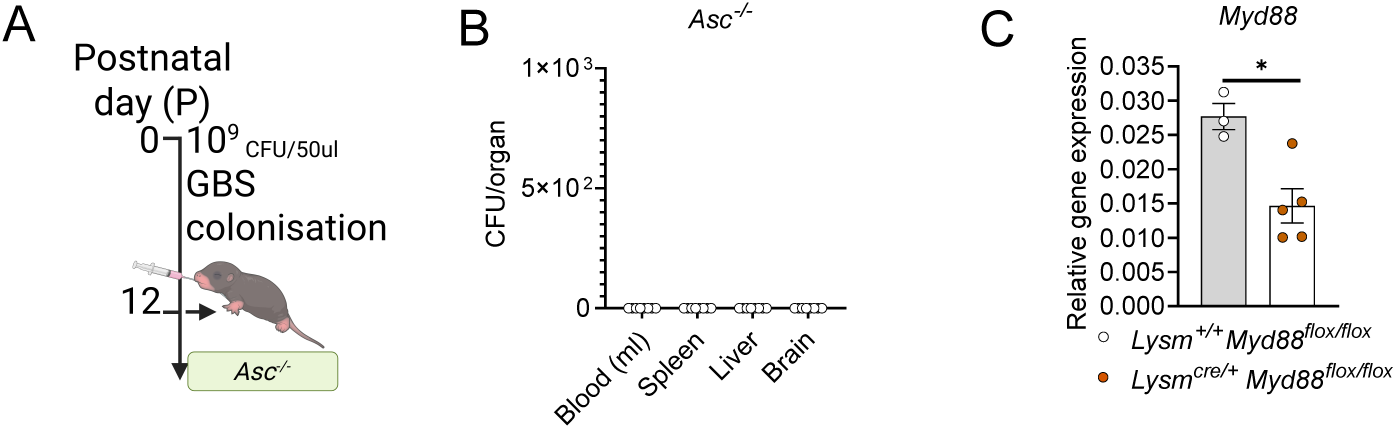
GBS colonization and downstream immune analyses. **A.** Experimental plan for colonization and analysis of *Asc^-/-^* at 2 days post-colonization at P14. **B.** CFU count of the GBS in the blood, Spleen, liver, and brain at 2 days post-colonization at P14. **C.** Quantitative real-time PCR (qRT-PCR) analysis of *Myd88* gene expression in Ly6C^-^MHC-II⁺ cells isolated from the colon of the *Lysm^cre/+^ Myd88^flox/flox^*(n=5), and *Lysm^+/+^ Myd88^flox/flox^* (n=3) mice.

## References

1. Seale, A. C. et al. Estimates of the burden of group B streptococcal disease worldwide for pregnant women, stillbirths, and children. Clin. Infect. Dis. 65, S200–S219 (2017).

2. Nanduri, S. A. et al. Epidemiology of invasive early-onset and late-onset group B streptococcal disease in the United States, 2006 to 2015: multistate laboratory and population-based surveillance. JAMA Pediatr. 173, 224–233 (2019).

3. Tazi, A. et al. Risk factors for infant colonization by hypervirulent CC17 group B Streptococcus: toward the understanding of late-onset disease. Clin. Infect. Dis. 69, 1740–1748 (2019).

4. Travier, L. et al. Neonatal susceptibility to meningitis results from the immaturity of epithelial barriers and gut microbiota. Cell Rep. 35, (2021).

5. Freudenhammer, M. et al. Invasive group B Streptococcus disease with recurrence and in multiples: towards a better understanding of GBS late-onset sepsis. Front. Immunol. 12, 617925 (2021).

6. Kolter, J. & Henneke, P. Codevelopment of microbiota and innate immunity and the risk for group B streptococcal disease. Front. Immunol. 8, 1497 (2017).

7. Tourneur, E. & Chassin, C. Neonatal Immune Adaptation of the Gut and Its Role during Infections. Clin. Dev. Immunol. 2013, 1–17 (2013).

8. Henneke, P. & Golenbock, D. T. Innate immune recognition of lipopolysaccharide by endothelial cells. Crit. Care Med. 30, S207–S213 (2002).

9. Mansoori Moghadam, Z., et al. Reactive oxygen species regulate early development of the intestinal macrophage-microbiome interface. Blood 145, 2025–2040 (2025).

10. Viola, M. F. & Boeckxstaens, G. Intestinal resident macrophages: Multitaskers of the gut. Neurogastroenterol. Motil. 32, e13843 (2020).

11. Bain, C. C. et al. Constant replenishment from circulating monocytes maintains the macrophage pool in the intestine of adult mice. Nat. Immunol. 15, 929–937 (2014).

12. Delfini, M., Stakenborg, N., Viola, M. F. & Boeckxstaens, G. Macrophages in the gut: masters in multitasking. Immunity 55, 1530–1548 (2022).

13. Hegarty, L. M., Jones, G.-R. & Bain, C. C. Macrophages in intestinal homeostasis and inflammatory bowel disease. Nat. Rev. Gastroenterol. Hepatol. 20, 538–553 (2023).

14. Kang, B. et al. Commensal microbiota drive the functional diversification of colon macrophages. Mucosal Immunol. 13, 216–229 (2020).

15. Chung, H. et al. Gut immune maturation depends on colonization with a host-specific microbiota. Cell 149, 1578–1593 (2012).

16. Grainger, J. R., Konkel, J. E., Zangerle-Murray, T. & Shaw, T. N. Macrophages in gastrointestinal homeostasis and inflammation. Pflüg. Arch. - Eur. J. Physiol. 469, 527–539 (2017).

17. Bain, C. C. & Schridde, A. Origin, differentiation, and function of intestinal macrophages. Front. Immunol. 9, 2733 (2018).

18. Serbina, N. V. & Pamer, E. G. Monocyte emigration from bone marrow during bacterial infection requires signals mediated by chemokine receptor CCR2. Nat. Immunol. 7, 311–317 (2006).

19. Tsou, C.-L. et al. Critical roles for CCR2 and MCP-3 in monocyte mobilization from bone marrow and recruitment to inflammatory sites. J. Clin. Invest. 117, 902–909 (2007).

20. Sbarbati, R. Morphogenesis of the intestinal villi of the mouse embryo: chance and spatial necessity. J. Anat. 135, 477 (1982).

21. Hopton, R. E., Jahahn, N. J. & Zemper, A. E. *Lrig1* drives cryptogenesis and restrains proliferation during colon development. Am. J. Physiol.-Gastrointest. Liver Physiol. 325, G570–G581 (2023).

22. Yip, J. L., Balasuriya, G. K., Spencer, S. J. & Hill-Yardin, E. L. The role of intestinal macrophages in gastrointestinal homeostasis: heterogeneity and implications in disease. Cell. Mol. Gastroenterol. Hepatol. 12, 1701–1718 (2021).

23. Isidro, R. A. & Appleyard, C. B. Colonic macrophage polarization in homeostasis, inflammation, and cancer. Am. J. Physiol.-Gastrointest. Liver Physiol. 311, G59–G73 (2016).

24. Zigmond, E. et al. Ly6Chi monocytes in the inflamed colon give rise to proinflammatory effector cells and migratory antigen-presenting cells. Immunity 37, 1076–1090 (2012).

25. Wright, P. B. et al. The mannose receptor (CD206) identifies a population of colonic macrophages in health and inflammatory bowel disease. Sci. Rep. 11, 19616 (2021).

26. Ma, S., Zhu, K., Liu, Y. & Wang, J. Research Progress on the Relationship Between the Intestinal Barrier and Macrophages. Curr. Issues Mol. Biol. 47, 813 (2025).

27. Huang, H. et al. Kupffer cell programming by maternal obesity triggers fatty liver disease. Nature 1–9 (2025).

28. Mass, E. et al. Specification of tissue-resident macrophages during organogenesis. Science 353, aaf4238 (2016).

29. Liu, Z. et al. Fate mapping via Ms4a3-expression history traces monocyte-derived cells. Cell 178, 1509–1525 (2019).

30. De Schepper, S. et al. Self-maintaining gut macrophages are essential for intestinal homeostasis. Cell 175, 400–415 (2018).

31. Priya, S. & Blekhman, R. Population dynamics of the human gut microbiome: change is the only constant. Genome Biol. 20, 150 (2019).

32. Lkhagva, E. et al. The regional diversity of gut microbiome along the GI tract of male C57BL/6 mice. BMC Microbiol. 21, 44 (2021).

33. Vadevoo, S. M. P., et al. The macrophage odorant receptor Olfr78 mediates the lactate-induced M2 phenotype of tumor-associated macrophages. Proc. Natl. Acad. Sci. 118, e2102434118 (2021).

34. Orecchioni, M. et al. Olfactory receptor 2 in vascular macrophages drives atherosclerosis by NLRP3-dependent IL-1 production. Science 375, 214–221 (2022).

35. Di Cara, F. et al. Peroxisomes in immune response and inflammation. Int. J. Mol. Sci. 20, 3877 (2019).

36. Meghnem, D., Leong, E., Pinelli, M., Marshall, J. S. & Di Cara, F. Peroxisomes regulate cellular free fatty acids to modulate mast cell TLR2, TLR4, and IgE-mediated activation. Front. Cell Dev. Biol. 10, 856243 (2022).

37. Yu, W. et al. Stat2-Drp1 mediated mitochondrial mass increase is necessary for pro-inflammatory differentiation of macrophages. Redox Biol. 37, 101761 (2020).

38. Su, J. et al. Structure of the intact Tom20 receptor in the human translocase of the outer membrane complex. PNAS Nexus 3, pgae269 (2024).

39. Iliev, I. D. et al. Microbiota-mediated mechanisms of mucosal immunity across the lifespan. Nat. Immunol. 1–15 (2025).

40. Bain, C. C. & Mowat, A. McI. Macrophages in intestinal homeostasis and inflammation. Immunol. Rev. 260, 102–117 (2014).

41. Russell, N. J. et al. Risk of early-onset neonatal group B streptococcal disease with maternal colonization worldwide: systematic review and meta-analyses. Clin. Infect. Dis. 65, S152–S159 (2017).

42. Carl, M. A. et al. Sepsis from the gut: the enteric habitat of bacteria that cause late-onset neonatal bloodstream infections. Clin. Infect. Dis. 58, 1211–1218 (2014).

43. Tazi, A. et al. Risk factors for infant colonization by hypervirulent CC17 group B Streptococcus: toward the understanding of late-onset disease. Clin. Infect. Dis. 69, 1740–1748 (2019).

44. Karampatsas, K. et al. Clinical risk factors associated with late-onset invasive group B streptococcal disease: systematic review and meta-analyses. Clin. Infect. Dis. 75, 1255–1264 (2022).

45. Berardi, A. et al. Group B streptococcal colonization in 160 mother-baby pairs: a prospective cohort study. J. Pediatr. 163, 1099–1104 (2013).

46. Joubrel, C. et al. Group B Streptococcus neonatal invasive infections, France 2007–2012. Clin. Microbiol. Infect. 21, 910–916 (2015).

47. Krause, J. C. et al. Very late-onset group B Streptococcus meningitis, sepsis, and systemic shigellosis due to interleukin-1 receptor-associated kinase-4 deficiency. Clin. Infect. Dis. 49, 1393–1396 (2009).

48. Yazji, I. et al. Endothelial TLR4 activation impairs intestinal microcirculatory perfusion in necrotizing enterocolitis via eNOS–NO–nitrite signaling. Proc. Natl. Acad. Sci. 110, 9451–9456 (2013).

49. Kordy, K. et al. Contributions to human breast milk microbiome and enteromammary transfer of Bifidobacterium breve. PloS One 15, e0219633 (2020).

50. Elling, R. et al. Synchronous recurrence of group B streptococcal late-onset sepsis in twins. Pediatrics 133, e1388–e1391 (2014).

51. Al Nabhani, Z. et al. A weaning reaction to microbiota is required for resistance to immunopathologies in the adult. Immunity 50, 1276–1288 (2019).

52. Maeda, K. et al. Wnt5a-Ror2 signaling between osteoblast-lineage cells and osteoclast precursors enhances osteoclastogenesis. Nat. Med. 18, 405–412 (2012).

53. Srinivas, S. et al. Cre reporter strains produced by targeted insertion of EYFP and ECFP into the ROSA26 locus. BMC Dev. Biol. 1, 4 (2001).

54. Daigle, T. L. et al. A suite of transgenic driver and reporter mouse lines with enhanced brain-cell-type targeting and functionality. Cell 174, 465–480 (2018).

55. Foley, J. W. et al. Gene expression profiling of single cells from archival tissue with laser-capture microdissection and Smart-3SEQ. Genome Res. 29, 1816–1825 (2019).

56. Bhardwaj, V. et al. snakePipes: facilitating flexible, scalable and integrative epigenomic analysis. Bioinformatics 35, 4757–4759 (2019).

57. Martin, M. Cutadapt removes adapter sequences from high-throughput sequencing reads. EMBnet J. 17, 10–12 (2011).

58. Dobin, A. et al. STAR: ultrafast universal RNA-seq aligner. Bioinformatics 29, 15–21 (2013).

59. Liao, Y., Smyth, G. K. & Shi, W. featureCounts: an efficient general purpose program for assigning sequence reads to genomic features. Bioinformatics 30, 923–930 (2014).

60. Martin, F. J. et al. Ensembl 2023. Nucleic Acids Res. 51, D933–D941 (2023).

61. Love, M. I., Huber, W. & Anders, S. Moderated estimation of fold change and dispersion for RNA-seq data with DESeq2. Genome Biol. 15, 550 (2014).

62. Smith, G. K. Limma: linear models for microarray data. Bioinformatics and Computational Biology Solutions using R and Bioconductor. Springer N. Y. 397–420 (2005).

63. Schubert, M. et al. Perturbation-response genes reveal signaling footprints in cancer gene expression. Nat. Commun. 9, 20 (2018).

64. Badia-i-Mompel, P., et al. decoupleR: ensemble of computational methods to infer biological activities from omics data. Bioinforma. Adv. 2, vbac016 (2022).

65. Godon, J. J., Zumstein, E., Dabert, P., Habouzit, F. & Moletta, R. Molecular microbial diversity of an anaerobic digestor as determined by small-subunit rDNA sequence analysis. Appl. Environ. Microbiol. 63, 2802–2813 (1997).

66. Klindworth, A. et al. Evaluation of general 16S ribosomal RNA gene PCR primers for classical and next-generation sequencing-based diversity studies. Nucleic Acids Res. 41, e1–e1 (2013).

67. Berry, D., Ben Mahfoudh, K., Wagner, M. & Loy, A. Barcoded Primers Used in Multiplex Amplicon Pyrosequencing Bias Amplification. Appl. Environ. Microbiol. 77, 7846–7849 (2011).

68. Lagkouvardos, I. et al. IMNGS: a comprehensive open resource of processed 16S rRNA microbial profiles for ecology and diversity studies. Sci. Rep. 6, 33721 (2016).

69. Edgar, R. C. UNOISE2: improved error-correction for Illumina 16S and ITS amplicon sequencing. BioRxiv 081257 (2016).

70. Edgar, R. C. UPARSE: highly accurate OTU sequences from microbial amplicon reads. Nat. Methods 10, 996–998 (2013).

71. Pruesse, E., Peplies, J. & Glöckner, F. O. SINA: accurate high-throughput multiple sequence alignment of ribosomal RNA genes. Bioinformatics 28, 1823–1829 (2012).

72. Quast, C. et al. The SILVA ribosomal RNA gene database project: improved data processing and web-based tools. Nucleic Acids Res. 41, D590–D596 (2012).

73. Lagkouvardos, I., Fischer, S., Kumar, N. & Clavel, T. Rhea: a transparent and modular R pipeline for microbial profiling based on 16S rRNA gene amplicons. PeerJ 5, e2836 (2017).

74. Jost, L. PARTITIONING DIVERSITY INTO INDEPENDENT ALPHA AND BETA COMPONENTS. Ecology 88, 2427–2439 (2007).

75. Chen, J. et al. Associating microbiome composition with environmental covariates using generalized UniFrac distances. Bioinformatics 28, 2106–2113 (2012).

